# Sequencing and comparative analysis of three *Chlorella* genomes provide insights into strain-specific adaptation to wastewater

**DOI:** 10.1101/630145

**Authors:** Tian Wu, Linzhou Li, Xiaosen Jiang, Yong Yang, Yanzi Song, Liang Chen, Xun Xu, Yue Shen, Ying Gu

## Abstract

Microalgal *Chlorella* has been demonstrated to process wastewater efficiently from piggery industry, yet optimization through genetic engineering of such a bio-treatment is currently challenging, largely due to the limited data and knowledge in genomics. In this study, we first investigated the differential growth rates among three wastewater-processing *Chlorella* strains: *Chlorella sorokiniana* BD09, *Chlorella sorokiniana* BD08 and *Chlorella sp.* Dachan, and the previously published *Chlorella sorokiniana* UTEX 1602, showing us that BD09 maintains the best tolerance in synthetic wastewater. We then performed genome sequencing and analysis, resulting in a high-quality assembly for each genome with scaffold N50 >2 Mb and genomic completeness ≥ 91%, as well as genome annotation with 9,668, 10,240, 9,821 high-confidence gene models predicted for BD09, BD08, and Dachan, respectively. Comparative genomics study unravels that metabolic pathways, which are involved in nitrogen and phosphorus assimilation, were enriched in the faster-growing strains. We found that gene structural variation and genomic rearrangement might contribute to differential capabilities in wastewater tolerance among the strains, as indicated by gene copy number variation, domain reshuffling of orthologs involved, as well as a ∼1 Mb-length chromosomal inversion we observed in BD08 and Dachan. In addition, we speculated that an associated bacterium, *Microbacterium chocolatum*, which was identified within Dachan, play a possible role in synergizing nutrient removal. Our three newly sequenced *Chlorella* genomes provide a fundamental foundation to understand the molecular basis of abiotic stress tolerance in wastewater treatment, which is essential for future genetic engineering and strain improvement.

## Introduction

Increasing demand on meat consumption has triggered rapid growth of the piggery industry throughout the world, especially in China ^1^, which is accompanied by increasing wastewater production and potential environmental problems. Excessive organic and inorganic pollutants containing nitrogen and phosphorus in wastewater without proper treatment often lead to eutrophication, oxygen depletion and ammonia toxicity to aquatic ecosystem ^2^, therefore, wastewater management is a growing environmental challenge worldwide.

In recent years, several biological approaches using microalgae to remove biological wastes have gained application. Microalgae is a large and diverse group of unicellular, photosynthetic, and chlorophyll-containing micro-organisms. Like plants, microalgae transmit solar energy into chemical compounds with fixed-carbons for biomass accumulation. Previous studies also showed that some green microalgae can grow in wastewater rapidly and absorb compounds containing nitrogen and phosphorus from the environment. Importantly, the algal biomass produced through such a process can be further used as feedstock, biofuel production, and for a series of by-products, such as drugs, foods, nutrition, fertilizers, and animal/fish feed supplements. Altogether, wastewater treatment by microalgae appears to be efficient for both biomass recycling and environmental protection in the piggery industry ^3^.

Among various microalgae species, *Chlorella* (belonging to Chlorellaceae, Chlorellales, Trebouxiophyceae), a genus of green microalga with diameters varying from 5 to 10 μm, is nowadays widely used in wastewater treatment with several advantages: rapid growth, simple and low-cost cultivation requirements, and a proven capability for producing high-value bioproducts, biomass, and biodiesel ^4^. The great commercial potential of the *Chlorella* genus (hereafter refer to as *Chlorella*) is thus a driving force in the effort of optimizing the process flow for wastewater treatment and expanding the applications in bioproduct manufacturing. However, challenges are still ahead, one of which is that wastewater is a harsh environment for most microalgal strains due to nutrient imbalance, deficiency of certain essential trace elements, and the presence of toxic compounds ^3^. In fact, some *Chlorella* strains do show good tolerance of certain wastewater environment in nature ^5 6^, driving the industry to continuously seek better strains for harsher conditions, especially for extremely high nutrient imbalance and salt concentrations, and to satisfy a growing market need for more efficient production of high-value metabolites.

One way to achieve *Chlorella* optimization is through genetic engineering. This can be achieved by random mutagenesis followed by selection for certain target phenotypic traits or by overexpression of certain transgenes with desired functions. Recently, rapidly evolved gene editing techniques make it possible for rapid and precise genetic modification for plants or algae ^7^. In order to apply such an engineering approach in various industrial purposes, we need to have the complete genome sequences with gene annotation information for target gene selection or genome-wide screening. However, only a few nuclear genomes of *Chlorella* clade are currently available in the NCBI database, including *Chlorella variabilis* NC64A ^8^ (hereafter referred to as NC64A), *Micractinium conductrix* ^9^, *Chlorella sorokiniana* UTEX 1602 ^9^ (hereafter referred to as UTEX 1602), and *Auxenochlorella protothecoides* ^10^, none of which has been tested in industrial wastewater. Furthermore, no genomic comparison between various *Chlorella* strains has been done to associate a genomic variation with a target phenotype, such as organic molecule clearance and stress-tolerance. Such a lack of knowledge in the genetics of *Chlorella* necessitates more efforts to obtain genomic blueprints of desirable *Chlorella* strains, together with various experiments on wastewater treatment.

In this study, we generated three high-quality whole-genome assemblies of industrial *Chlorella* strains, including *Chlorella sp.* Dachan (hereafter referred to as Dachan), *Chlorella sorokiniana* BD09 and *Chlorella sorokiniana* BD08 (hereafter referred to as BD09 and BD08). The three high-quality assemblies were further annotated by combining homolog-based, *ab initio*, and transcriptome-based prediction approaches, resulting in high-confidence gene models with >90% of the genes assigned with putative functions. By comparative analyses across five *Chlorella* strains, we discovered a group of genes associated with nitrogen metabolism and endocytosis pathways and varied domains of nitrogen and phosphorus transporters that may contribute to nutrient clearance from wastewater. Interestingly, an associated bacterium was observed within Dachan, which had been shown to improve the nitrogen and phosphorus absorbing ability for algae^11^. A chromosomal inversion event was confirmed in the genomes of two industrial strains, which still awaits future test on its adaptation in wastewater tolerance. Altogether, our study provides a rich resource of genomic information for some strains of *Chlorella* genus and lays the foundation for genetic engineering of microalgae system to improve their capabilities for wastewater treatment and accompanied commercial values.

## 1 Results

### 1.1 Comparison of growth performance in synthetic wastewater among *Chlorella* strains

Our pervious field trials of BD09 showed good performance in piggery wastewater treatment, with an estimation of the clearance efficiency as 72%, 88% and 61% in chemical oxygen demand (COD), total nitrogen and total phosphorus, respectively (see Table S1 for details). The nitrogen and phosphorus elements are enriched in wastewater mainly in the form of NH_4_^+^ (ammonia), NO_2_^−^ (nitrite), NO_3_^−^ (nitrate) and PO_4_^3−^ (orthophosphate), which are essential nutrients for *Chlorella* growth but may lead to abiotic stress at high concentrations. In order to comprehensively compare the performance of different *Chlorella* respond to wastewater with rich carbon, nitrogen and phosphorus compounds in the lab, as well as to explore the underlying molecular mechanism responsible for this capability, we made one-to-one growth rate comparisons among *Chlorella* strains in synthetic wastewater with various concentrations of carbon, nitrogen and phosphorus. In addition to BD08, BD09 and Dachan, the strain *Chlorella sorokiniana* UTEX 1602, the genome draft of which was reported recently ^9^, was included into the growth assay as a control to facilitate further comparative genomic analyses (Figure 1).

**Figure 1.**
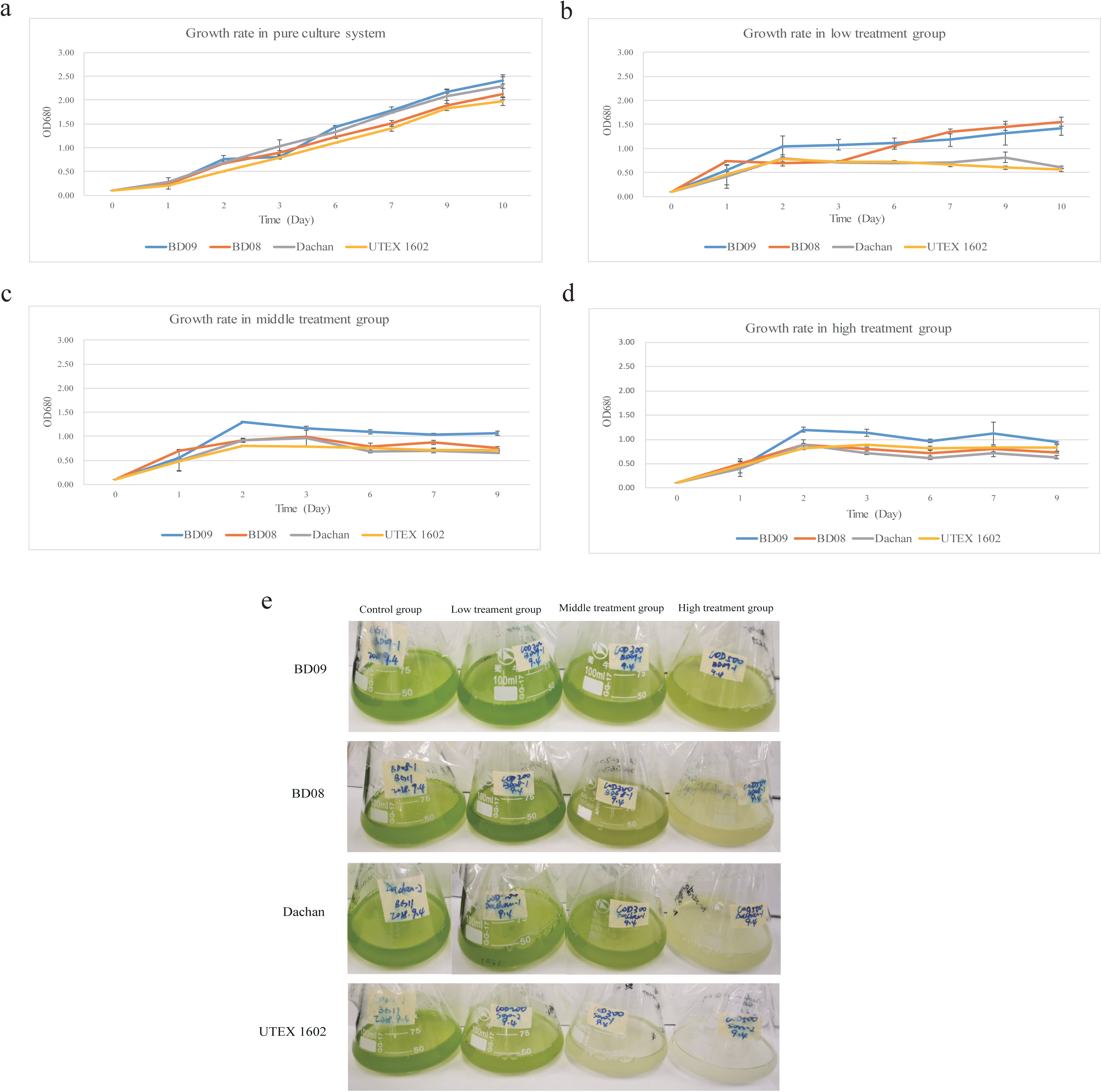
Experimental assessment of the tolerance capability in synthetic wastewater. Comparison of growth rates of four *Chlorella* strains (*Chlorella sorokiniana* BD09, *Chlorella sorokiniana* BD08, *Chlorella sp.* Dachan, and *Chlorella sorokiniana* UTEX 1602) in a) pure BG11 culture system as the control group, b) low concentration treatment group, c) middle concentration treatment group, and d) high concentration treatment group. The visualization of concentrations among the four strains in synthetic wastewater culture for 3 days was present in e).

All four *Chlorella* strains grew at comparable rate in the basic BG11 pure culture medium (Figure 1a), suggesting that their capacity of growth was similar without abiotic stress. However, when the concentrations of glucose, NH_4_Cl and KH_2_PO_4_ were increased to 200 mg/L, 59.43 mg/L and 8.78 mg/L, respectively, which were defined as the low concentration treatment group (see Methods), Dachan and UTEX 1602 failed to grow beyond day 6, while BD08 and BD09 were only mildly affected (Figure 1b, 1e). When the concentrations of glucose, nitrogen and phosphorus were further increased (middle and high concentration treatment group), the growth of BD09 was least affected in comparison with other three strains in the first few days, though none of the four strains could continue to grow beyond day 5 (Figure 1c, 1d, 1e), indicating that the concentration of carbon, nitrogen and phosphorus is a major factor to suppress the growth of these *Chlorella*, while BD09 appears to show the best resistance to such abiotic stress followed by BD08.

### 1.2 Genome sequencing and characteristics

In order to delineate the underlying molecular mechanism responsible for such a differential growth rate among the three *Chlorella* in wastewater, we sequenced the whole genomes of BD09, BD08, and Dachan using a whole-genome shotgun strategy on the HiSeq platform. In total, 43.16 Gb (799 X), 43.98 Gb (751 X), and 41.09 Gb (680 X) of clean data for BD09, BD08, and Dachan, respectively, were generated after data filtering (Table S2). K-mer frequency distribution analysis ^13^ was performed to estimate the genome complexity (see Methods). The 17-mer frequency distribution curve for the three newly sequenced genomes exhibited one-fold peak, indicates that all three genomes are simple genomes with low heterozygosity rate and simple repeat content (Figure S1). The estimated genome sizes of these three genomes were 55.1 Mb for BD09, 56.5 Mb for BD08, and 60.4 Mb for Dachan.

Genomes were assembled and optimized by Platanus ^14^, and then a series of evaluations was performed for assembly contiguity and completeness (see Methods). Three high-quality assemblies were finalized with a genome size of 54.0 Mb for BD09, 58.6 Mb for BD08, 60.4 Mb for Dachan, respectively, highly consistent with each predicted genome size by the 17 K-mer analysis ^13^. The scaffold N50 sizes were 3.58 Mb for BD09, 3.63 Mb for BD08 and 2.58 Mb for Dachan, and more than 94.5% of bases for BD09, 92% for BD08, and 90% for Dachan were located in scaffolds larger than 1 Mb, which were much higher than the other genomes of *Chlorella* clade reported previously (71.4% for NC64A, 88.3% for UTEX 1602, and 60% for *Micractinium conductrix* SAG 24.880, respectively) (Table 1, Table S3). The RNA-mapping rate and BUSCO ^15^ dataset support a high assembly completeness for all three genomes, suggesting 95.7%, 95.7%, 91% of the assemblies were covered by transcriptome evidences, and 91.8%, 91.6%, 88.7% of the Chlorophyta genes were aligned to our assembled sequences. We further assembled chloroplast and mitochondrial genomes for the newly sequenced genomes, obtaining chloroplast genome sequences of 131,724 bp for BD09, 109,940 bp for BD08, and 113,449 bp for Dachan, respectively. For mitochondrial genomes, the genome sizes were 57,540 bp, 53,743 bp, and 45,617 bp for BD09, BD08, and Dachan, respectively. All of these results strongly suggest that all our newly sequenced *Chlorella* genomes have a higher quality of genome assemblies both in contiguity and in completeness than other *Chlorella* clade genomes reported so far (Table 1, Table S3) ^8 9^.

**Table 1.** Summary of characteristics in genome assembly and annotation for *Chlorella sorokiniana* BD09, *Chlorella sorokiniana* BD08 and *Chlorella sp.* Dachan in comparison with other published *Chlorella* clade genomes.

|  | *Chlorella sorokiniana* BD09 | *Chlorella sorokiniana* BD08 | *Chlorella* sp. Dachan | *Chlorella variabilis* NC64A | *Chlorella sorokiniana* UTEX 1602 | Micractinium conductrix SAG 241.80 |
| --- | --- | --- | --- | --- | --- | --- |
| Genome size (Mb) | 54.0 | 58.6 | 60.4 | 46.2 | 59.6 | 61.0 |
| GC content | 65.3% | 63.8% | 65.1% | 67.0% | 63.9% | 67.1% |
| Contig N50 (bp) | 35,822 | 42,232 | 21,305 | 27,649 | 2,592,956 | 1,210,495 |
| Scaffold N50 (bp) | 3,584,288 | 3,632,890 | 2,575,675 | 1,469,606 | 2,592,956 | 1,210,495 |
| % of larger than 1 Mb | 94.5% | 92.0% | 90.0% | 71.4% | 88.3% | 60.0% |
| % of Ns gaps | 0.82% | 1.84% | 4.16% | 8.55% | 0.00% | 0.00% |
| % of RNA mapping | 95.70% | 95.70% | 91.00% | / | 95.03% | 94.57% |
| % of DNA mapping | 99.16% | 98.13% | 95.80% | / | 99.29% | 98.92% |
| # of gene models | 9,668 | 10,240 | 9,821 | 9,791 | 9,587 | 9,349 |
| Average protein length (aa) | 578 | 553 | 535 | 456 | 631 | 637 |
| # of exons/gene | 12.8 | 12.3 | 12.1 | 7.3 | 14.6 | 13.9 |
| Average exon length (bp) | 135 | 134 | 132 | 170 | 146 | 152 |
| Average intron length (bp) | 217 | 228 | 236 | 209 | 231 | 250 |
| Coding sequence (%) | 31.06% | 28.95% | 26.01% | 29.00% | 30.60% | 29.40% |

Note: Not including chloroplast, mitochondrial genomes or associated bacterial genomes.

Through structural annotation and comparison, we demonstrated that the newly sequenced *Chlorella* genomes all maintain a simple repeat composition, covering only 3%∼5% of the genome which were dominantly by LTRs (long terminal repeats) (Table S4). Gene annotation was performed by a combinatorial strategy integrating the homolog-based, *ab initio*, and transcriptome-aided approaches (see Methods), resulting in 9,668 protein-coding gene models for BD09, 10,240 for BD08, and 9,821 for Dachan, respectively (Table 1, Table S5). Evaluation in gene completeness shows that ∼90% of gene models could be assigned by at least one known functional protein databases. Furthermore, we found no significant differences in genic characteristics among the six *Chlorella* clade genomes, both in average numbers and length distribution of genes, exons, and introns (Table 1).

### 1.3 Phylogeny and gene families

A total of 362 single-copy gene families from fourteen genomes (Table S9, Methods) were identified by Orthofinder ^16^. These gene families were then used following a concatenation-based approach to reconstruct the phylogenetic tree of the *Chlorella* strains we sequenced together with their closely-related species. By implementing the CAT+GTR amino acid substitution model, we generated a maximum likelihood phylogenetic tree for the selected green algae, with the red alga *Cyanidioschyzon merolae* ^17^ as the outgroup. The results confirmed that all three strains belong to the *Chlorella* genus, with BD09 evolutionarily closest to UTEX 1602, followed by BD08 and Dachan (Figure 2b). We also compared the overall identity of amino-acid sequences of these 14 genomes (Figure S5). Consistently, BD09, BD08, Dachan, and UTEX 1602 shows the highest similarity, indicates that they diverged recently from the last common ancestor.

**Figure 2.**
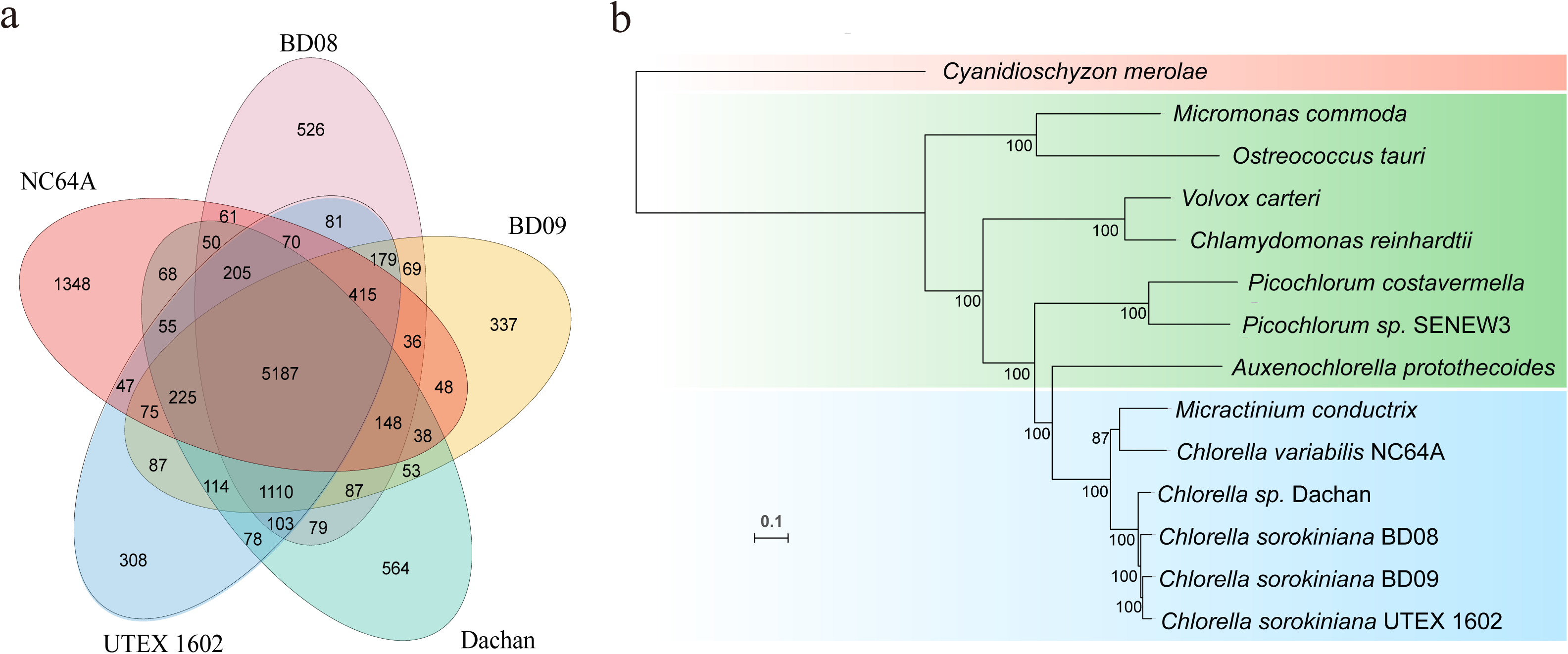
Gene family analysis and species phylogeny reconstruction. a) Venn diagram showing shared and unique orthogroups among *Chlorella strains: Chlorella sorokiniana* BD09, *Chlorella sorokiniana* BD08, *Chlorella sp.* Dachan, *Chlorella sorokiniana* UTEX 1602, and *Chlorella variabilis* NC64A. b) Phylogenetic tree reconstruction of the Chlorella genus with other closely-related algae, with *Cyanidioschyzon merolae* as the outgroup. A total of 362 single-copy gene families were extracted from the fourteen algal genomes, followed by multiple sequence alignment of the protein sequences by MAFFT. The alignments were refined by filtering out ‘poor regions’ with >50% of the orthologous sites represented as ‘N’ by Gblocks. The final alignments for each species were further concatenated into a supermatrix for the reconstruction of a maximum likelihood phylogenetic tree by RAxML, with CAT+GTR amino acid substitution model.

To identify genetic variations that may link to distinct physiological features of the *Chlorella* strains, we performed gene family analysis. Both shared and species-specific gene families among five closely-related *Chlorella* strains (BD09, BD08, Dachan, UTEX 1602, and NC64A) were analyzed according to the orthogroups generated by Orthofinder (Figure 2a). Within all 8,804 orthogoups, 7,874 were present in BD09, of which 7,231, 6,962, 7,392, and 6,172 genes were shared with BD08, Dachan, UTEX 1602, and NC64A, respectively. Unlike BD09, NC64A is not an industrial strain known for wastewater treatment, we then in particular compared gene families between BD09 and NC64A and found 1,702 expanded gene families and 573 contracted gene families in the BD09 genome. Interestingly, for these expanded gene families identified in BD09, they were significantly enriched in metabolism-related pathways as indicated in our KEGG pathway enrichment analysis, including pathways that function in endocytosis, starch and sucrose metabolism, and glycan degradation (Table S6). As around three-quarters of the organic carbon in wastewater is present as carbohydrates, fats, proteins, amino acids, and volatile acids, expansion of these metabolic gene families may contribute to the better growth of BD09 in wastewater.

We then expanded our comparison analysis in gene families between BD09, BD08, and Dachan with that in NC64A to identify a set of well-supported gene families specific in the industrial strains to explain their possible adaptive capabilities to industrial wastewater environment. In total, 3,467 gene families (7,305 genes) were totally absent in the NC64A genome, but present in at least one of the three industrial strain genomes (Figure 2a).Consistently, we found that 3,467 gene families were functional enriched in metabolic pathways in our KEGG analysis, including endocytosis, nitrogen metabolism, glycan degradation, phosphonate and phosphinate metabolism, and starch and sucrose metabolism (Table S7). In sum, these results imply that gene copy number variation in a couple of gene families involved in nutrition (C/N/P) absorption and metabolism may contribute to the better performance of our newly sequenced *Chlorella* strains in wastewater.

### 1.4 Comparative genomics and collinear blocks

In order to further characterize the genomic differences between our newly sequenced *Chlorella* genomes and the other algal strains, we conducted synteny and collinearity analysis between genomes by MCScanx ^18^. Circos ^19^ was used to visualize gene density, TE composition, GC content, and scaffold-scale homologous relationships between the three new genomes and the genome of UTEX 1602. The result showed a high degree of collinearity between genome pairs, with more than 800 collinear blocks detected for each pair (Figure 3a, Figure 4a-4c, Figure S3-S4). Our genome-wide comparison consolidates the conclusion that BD08 and BD09, which are cultivated in wastewater, are both derived from laboratory *Chlorella sorokiniana* SU1. Interestingly, Dachan, with an unknown origin, maintains a high degree of similarity with UTEX 1602, indicating a recent divergence of these two strains (Figure 3a).

**Figure 3.**
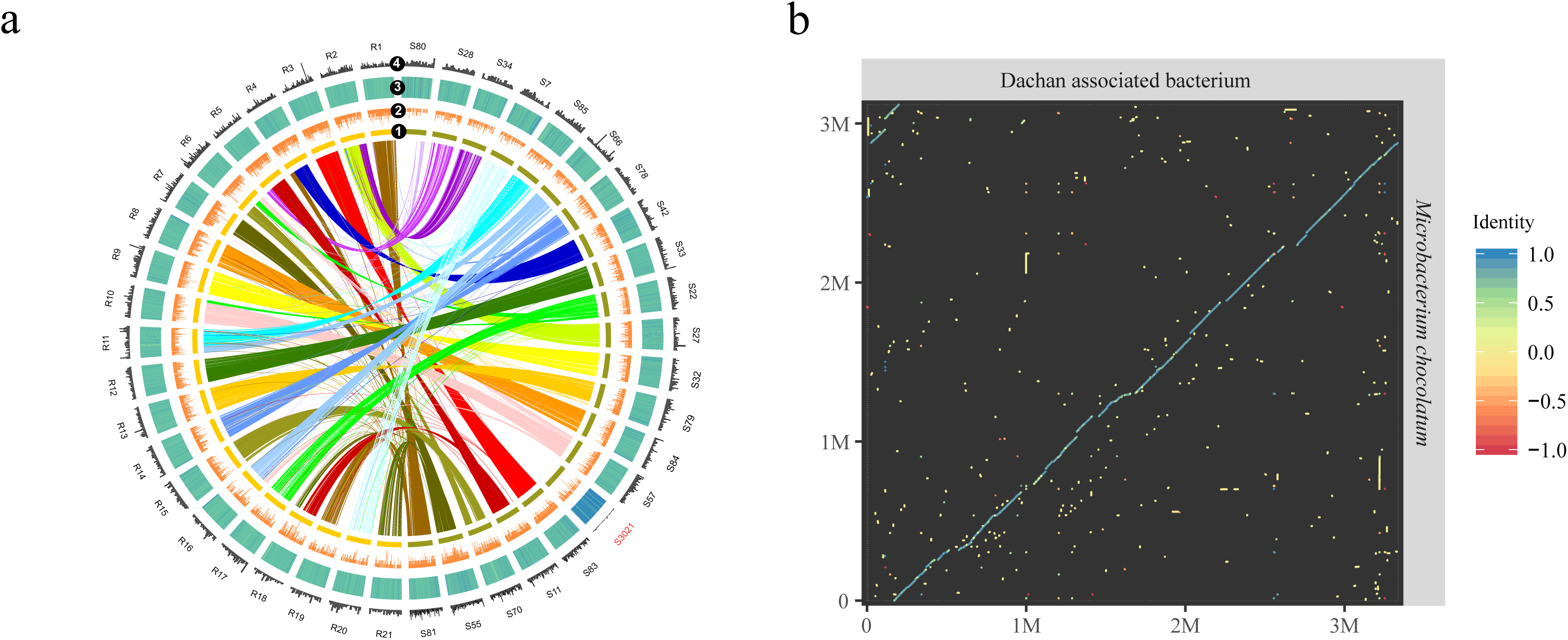
Comparison and validation of an associate bacterium identified in Dachan by synteny analysis. a) Circos visualization of collinearity between *Chlorella sp.* Dachan (S) and *Chlorella sorokiniana* UTEX 1602 (R), only scaffolds with length >1Mb were used. Different layers denoted: 1) collinear regions between the two strains connected by colored lines, 2) gene density, 3) GC content, and 4) TE density. b) Dot-plot visualization of the alignment between the associated bacterium identified in the *Chlorella sp.* Dachan genome and the published bacterium: *Microbacterium chocolatum*.

**Figure 4.**
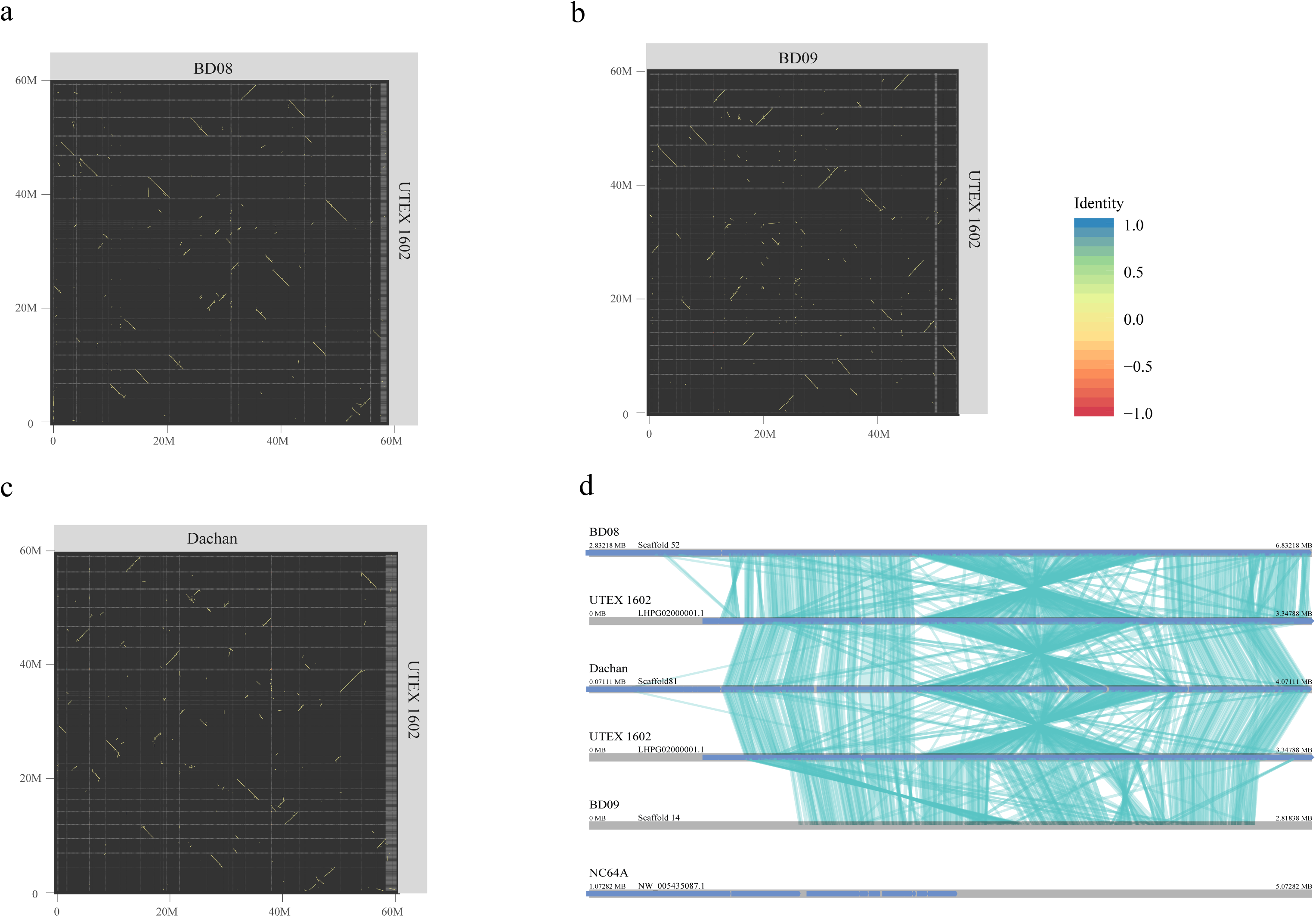
Genome-wide syntenic/collinear blocks and a chromosomal inversion event. Dot-plot visualization of collinearity between a) *Chlorella sorokiniana* BD08 and *Chlorella sorokiniana* UTEX 1602, between b) *Chlorella sorokiniana* BD09 and *Chlorella sorokiniana* UTEX 1602, between c) *Chlorella sp.* Dachan and *Chlorella sorokiniana* UTEX 1602. d) A chromosomal inversion event was identified in *Chlorella sorokiniana* BD08 and *Chlorella sp.* Dachan, in comparisons with *Chlorella sorokiniana* UTEX 1602 and *Chlorella sorokiniana* BD09, respectively. This corresponding region of that scaffold was absent in *Chlorella variabilis* NC64A.

It is worthwhile to notice that Dachan performs well in our field trials of piggery wastewater treatment, yet grows poorly in synthetic wastewater, leaving us a puzzling phenomenon in Dachan that calls for detailed genome-level analysis. Remarkably, we found a scaffold in the Dachan genome with totally different GC content and TE density compared with other scaffolds (Figure 3a). Moreover, this scaffold did not share any synteny blocks with the other algal genomes (Figure 3a), suggesting a distinct origin. We aligned this scaffold against the NCBI NR (non-redundant) database and found a good alignment with the genome sequence of a bacterium *Microbacterium chocolatum*, as shown by the dot-plot analysis (Figure 3b). In order to test if this was due to bacterial contamination, we calculated the reads coverage between this scaffold and the other scaffolds of the Dachan genome, yet found little difference in coverage depth (Figure S2). The result indicates that the bacterium genome sequences in the Dachan strain might come from symbiosis. Whether such symbiosis, if any, may contribute to the adaptation of *Chlorella* to certain wastewater conditions needs further investigation.

To further investigate the genomic structural variations of these *Chlorella* strains, we used UTEX 1602 as the reference and confirmed that a genomic region of approximately 1 Mb in length (∼2% of the whole genome) had undergone a chromosome inversion in both BD08 and Dachan. Although BD08 is phylogenetically more related to BD09 than to Dachan, we did not see this inversion in BD09. These data suggested a possible genomic rearrangement hotspot and the inversion occurred independently in BD08 and Dachan (Figure 4d). We then conducted KEGG enrichment analysis for genes located in this rearrangement hotspot region, showing that genes were enriched in DNA replication, spliceosome, base excision repair, and homologous recombination, thus indicating this region is important for maintaining normal DNA replication and the genome integrity (Table S8). Interestingly, NC64A, which has not been reported for use in wastewater, did not contain this synteny region (Figure 4d). More experiments are required in the future to measure the expression level of genes in this region, as well as the maintenance capability of genome stability under various stress conditions to explore the possible biological significance of this rearrangement event.

### 1.5 Genes in nitrogen and phosphorus assimilation, transportation and regulation

Wastewater in piggery industry often contains a high concentration of nitrogen and phosphorus, which need to be removed before wastewater can be discharged into the environment. To gain molecular insights into the varied capabilities of certain *Chlorella* strains to tolerate high levels of nitrogen and phosphorus (Figure 1), we surveyed the genes involved in the assimilation and transport of nitrogen and phosphorus across six green algal genomes (BD09, BD08, Dachan, UTEX 1602, NC64A, and *Ostreococcus tauri* ^20^) (Table 2, Figure 5). Table 2 lists the copy numbers of genes involved in nitrogen assimilation and its regulation based on a target-searching approach (see Methods). For the nitrate assimilation pathway, which includes two steps both in transport and reduction, we observed a similar pattern in gene copy numbers and gene structural conservation across the five *Chlorella* strains (Table 2), but except for *Ostreococcus tauri*, the later experienced a contraction in the related genes. The genes involved in the transformation from nitrate to amino acid in the nitrate assimilation pathway were highly conserved in gene structure, but varied in copy numbers between different algae (Table 2). We found that nitrate transporter NRT1 was present in all five *Chlorella* strains, yet absent in *Ostreococcus tauri*. The nitrate transporter NRT2 proteins have two forms, one only contained the MFS superfamily (Major Facilitator Superfamily), the other one was a two-component system, which required a second protein, high-affinity nitrate assimilation related protein (NAR2), to be fully functional. This two-component system was only discovered in the two *Chlorella* strains, BD09 and UTEX 1602. For nitrate reductase (NR), BD08 and UTEX 1602 had the largest copy numbers of genes (5 copies), followed by BD09 (3 copies), Dachan (3 copies), and NC64A (1 copy), which might facilitate nitrogen assimilation (Table 2). Noteworthily, NC64A maintained five copies of ammonium transporters gene (AMT), relative higher than that in the other five green algae, suggesting a possible differential adaptation strategy developed to assimilate nitrogen.

**Table 2.** Copy numbers of genes involved in nitrate assimilation, transportation and regulation in green algae.

| Activity | Reference<br>Gene<br>Name | *Chlorella*<br>*sorokiniana*<br>BD08 | *Chlorella*<br>*sorokiniana*<br>BD09 | *Chlorella*<br>*sp.*<br>Dachan | *Chlorella*<br>*sorokiniana*<br>UTEX 1602 | *Chlorella*<br>*variabilis*<br>NC64A | *Ostreococcus*<br>*tauri* |
| --- | --- | --- | --- | --- | --- | --- | --- |
| Nitrate transporter | NRT1/<br>PTR | 3 | 3 | 3 | 3 | 2 | 0 |
| Nitrate transporter | NRT2 | 2 | 2 | 2 | 2 | 2 | 1 |
| Nitrate reductase | NR | 5 | 3 | 3 | 5 | 1 | 3 |
| Nitrite reductase | NiR | 1 | 1 | 1 | 1 | 1 | 1 |
| Glutamine synthetase | GLN | 1 | 1 | 1 | 1 | 2 | 0 |
| Glutamate synthase,<br>NADH-dependent | GSN/GSF | 2 | 2 | 2 | 3 | 3 | 1 |
| Molybdate-anion transporter | MOT2 | 1 | 1 | 1 | 2 | 1 | 2 |
| Molybdate transporter | MOT1 | 0 | 0 | 0 | 0 | 1 | 1 |
| Molybdopterin cofactor<br>sulfurase family<br>protein | CNX | 5 | 5 | 5 | 6 | 6 | 5 |
| Ammonium transporter | Amt | 3 | 3 | 1 | 3 | 5 | 4 |

**Figure 5.**
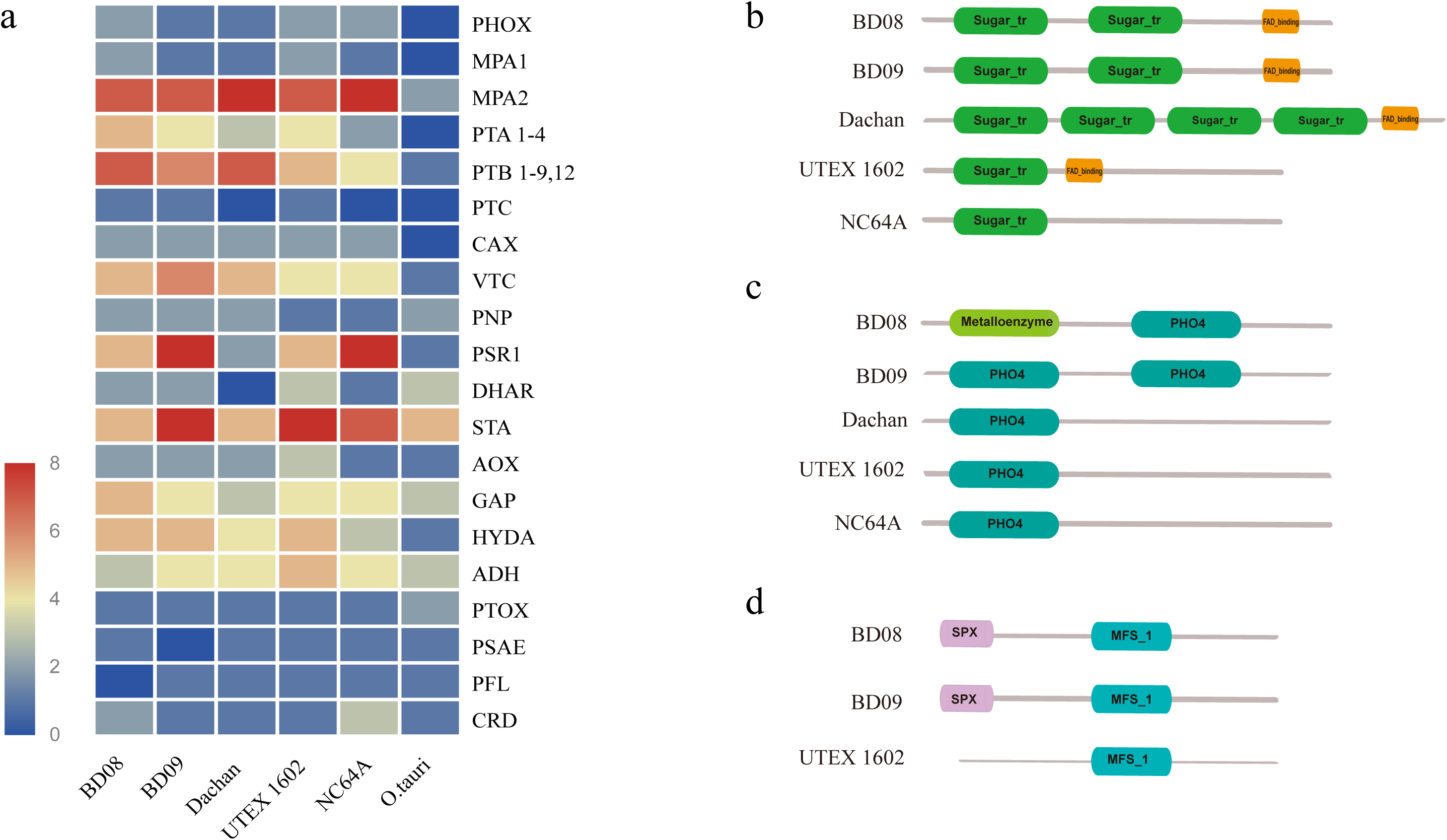
Analyses and comparison of gene families involved in the phosphorus assimilation pathway among *Chlorella sorokiniana* BD09, *Chlorella sorokiniana* BD08, *Chlorella sp.* Dachan, *Chlorella sorokiniana* UTEX 1602*, Chlorella variabilis* NC64A, and *Ostreococcus tauri*. a) Heatmap of genes related with phosphorus uptake and regulation. Domain analyses of three major phosphate transporters, b) domains of proton/phosphate symporter (PTA), c) domains of sodium/phosphate symporter (PTB), and d) domains of SPX domain-containing membrane protein (PTC) for different *Chlorella* strains.

Similar to the nitrogen assimilation analyses, we analyzed candidate genes involved in the transport and regulation of phosphorus across six green algae (Figure 5a). Genes responsible for phosphorus transporters, including proton/phosphate symporter (PTA), sodium/phosphate symporter (PTB), and SPX domain-containing membrane protein (PTC) were expanded in BD08 (5 copies for PTA, 7 copies for PTB, and 1 copy for PTC) than that in NC64A (2 copies for PTA, 4 copies for PTB, and no gene for PTC). Interestingly, we found that the domain gains and losses within the phosphorus transporter genes between different green algae (Figure 5b-5d). For instance, the PTA gene appeared to lose a flavin adenine dinucleotide (FAD) binding domain in NC64A compared with the other four *Chlorella* genus strains, which may compromise the protein function and lead to reduced ability for phosphorus assimilation (Figure 5b). The PTB gene in BD08 and BD09 had two phosphorus transporter family (PHO4) domains, while UTEX 1602, Dachan, and NC64A contained only one PHO4 domain (Figure 5c). Similarly, the SPX domain in PTC gene (Figure 5d) was present in both BD08 and BD09, but absent in UTEX 1602. In contrast, the whole PTC protein was absent in the other three algae, Dachan, NC64A, and *Ostreococcus tauri*.

Taken together, these data suggest that gene copy number variation and domain structure changes may contribute to the strain-specific capability in dealing with abiotic stress in wastewater, thus providing a clue for target functional testing and genetic modification, which eventually could help us to screen candidate microalgal strains with efficient capabilities in wastewater treatment and biomass production.

## 2 Discussion

In this study, the strains BD09 and BD08 were domesticated from *Chlorella sorokiniana* SU1, both showing good capability of tolerance in synthetic wastewater. The strain Dachan was originally collected from sewage and was assumed to have better performance in wastewater culture. However, we did not observe a higher growth rate of Dachan compared with the commercial strain UETX 1602 in the entire process of the synthetic wastewater experiments, and did find a much lower growing rate than that in BD09 at later treatment stages. Given that genomic analyses unraveled an associated bacterial genome in the Dachan genomic data, we assumed that the unexpected lower wastewater tolerance of Dachan was probably due to an unintentional removal of the associated bacterium by bacterial antibiotics, the latter were usually used in the treatment of synthetic wastewater, to avoid the bacterial contamination and disruption of OD_680_ measurement. Future work is needed to make it clear of the role of the associated bacterium identified in the Dachan genome in this study.

An extensive comparative genomics analysis was conducted between our three newly sequenced strains, which have been routinely used for industrial wastewater treatment, with the genome of NC64A strain that lacks the ability of growing in wastewater. We found that specific gene families identified in our industrial strains were significantly enriched in metabolic pathways, including endocytosis, nitrogen metabolism, glycan degradation, phosphonate and phosphinate metabolism, and starch and sucrose metabolism. Endocytosis, an essential process for cells to uptake extracellular nutrition, plays a key role to balance environmental responses and intracellular metabolites. Previous reports revealed that endocytosis is involved in the coordination of transportation of phosphorus and nitrogen transportation by several mechanisms. For instance, endocytosis plays a critical regulatory role in the degradation of plasma membrane proteins, including plant phosphate transporter 1 (PHT1) ^21 22^. Clustering of a high affinity nitrate transporter in endocytosis has also been reported to help avoid accumulation of ammonium to the toxic level under high ammonium stress condition ^23^. The enriched metabolic pathways in nitrogen, sucrose, and glycan detected in BD09 strain (Table S6, Table S7) were consistent with the observation of its superior growth performance in our synthetic wastewater system (Figure 1), providing an underlying mechanism that enables BD09 a better performance in nutrient removal and stress-tolerance compared with the other strains.

To investigate the dynamics of gene families involved in the nitrogen assimilation pathway, we included additional four *Chlorella* genomes for comparison in our study, besides of NC64A and *Ostreococcus tauri*, that was present in Emanuel Sanz-Luque et.al ^24^. To filter out false positives and retrieve false negatives in the identification of gene homologs, we implemented much stricter criteria and the most updated versions of genome dataset, compared with Emanuel Sanz-Luque’s study, in each step of our searching pipeline (see Methods). We found fewer copies of genes (NRT1, GLN, and CNX) were confirmed in this study than that reported previously in the genomes of NC64A and *Ostreococcus tauri*. On the other hand, we also found that NC64A had the largest number of genes in ammonium transporters (5 copies) among all the green algal genomes in comparison (Table 2), which was not present in Emanuel Sanz-Luque’s study ^24^. This observation provides new evidence to support that NC64A develops an alternative adaptation strategy, not using nitrate or nitrite but instead ammonium or amino acids to acquire nitrogen ^24 25^.

Further, we analyzed gains and losses of genes and the diversification of the structural domains across strains. NAR2, for example, a high-affinity nitrate assimilation related protein that interacts with protein NRT2, was only found in BD09 and UTEX 1602, which might explain an improved capability in nitrate absorption. For phosphate transporter genes, we found that the PTC protein was totally absent in Dachan, NC64A, and *Ostreococcus tauri*, while for the other algae, a conserved SPX domain of PTC genes were only present in BD08 and BD09, but absent in UTEX 1602. Recently, several SPX domain-containing proteins from the SPX subfamily have been confirmed to play a role in phosphate sensing, signaling, transport, storage, and metabolism, enabling plants to maintain phosphate homeostasis ^26 27 28^. We thus concluded that the SPX domain may play a critical role in phosphorus assimilation for BD08 and BD09, and contribute to the superior tolerance to synthetic wastewater in high concentrations of phosphorus. These findings open a door for further experiment to measure the capability in the absorbance and metabolism of phosphorus in these *Chlorella* strains.

A symbiotic association with the bacterium *Microbacterium* in *Chlorella* is a common phenomenon in nature. A bacterial symbiont G+C gram-positive bacterium CSSB-3 was previously isolated from *Chlorella sorokiniana.* It shares 98.6% similarity of 16S rRNA sequence with that of *Microbacterium trichotecenolytcum* and has been found to promote the growth of *Chlorella* ^29^. Another example is a symbiotic bacterium *Microbacterium sp.* HJ1, isolated from *Chlorella vulgaris*, shows 99% identity with a wastewater isolated bacterium - *Microbacterium kitamiense*. This bacterium has been proven to promote *Chlorella* growth in the modified livestock wastewater ^30^. Interestingly, we assembled a complete genome of the associated bacterium *Microbacterium chocolatum* that also belongs to *Microbacterium* genus in Dachan. *Microbacterium chocolatum* and *Microbacterium kitamiense* shares an identity over 99% as measured by 16S rRNA genes ^31^. In contrast, we did not observe an associated or symbiotic bacterium in BD08 or BD09. The possible reason was that Dachan was originally isolated from sewage, and adaptively cultured in wastewater or BG11 without sterilization, while both BD08 and BD09 were originated from a lab strain after long time culture which requires a bacterial free environment. When the associated bacterium was removed from our synthetic wastewater by antibiotics treatment, the wastewater tolerance of Dachan was dramatically reduced, while that of BD08 and BD09 remains the same, suggesting that diverse strategies and mechanisms may exist for *Chlorella* to grow in industrial environments.

Chromosomal inversion events were observed at the same region in both BD08 and Dachan, two independent strains with different origins. Such an inversion event was not found in BD09 or UTEX 1602, although they share the same origin with BD08, suggesting that the inversion region may be a rearrangement hotspot, and has been acquired independently by BD08 and Dachan. More interestingly, this 1 Mb region was totally absent in NC64A, the strain that was much less tolerant to industrial wastewater culture, posing a possibility that the structural alteration of this region is related to stress tolerance (Figure 4). Further analyses of genes in this inversion region indicated an enrichment of pathways involved in DNA repair and homologous recombination (Table S8). Further experiments on the expression levels of genes in this region, as well as the maintenance of genome stability under various stress conditions, will be interesting to conduct in order to determine the biological significance of this genomic rearrangement event.

Overall, our comparative genomics analysis provides some clues to associate the potential genotype with phenotypes of interest in our wastewater-tolerance *Chlorella* experiment. We found that some metabolic pathways such as nitrogen and phosphorus metabolism were enriched in the newly sequenced industrial strains, indicating a potential molecular basis for their tolerance in wastewater. Investigation of genes related with nitrogen and phosphorus assimilation provides insights into the underlying molecular mechanism responsible for nutrient removal in wastewater. Furthermore, a chromosomal inversion event and associated bacterium, might also contribute to the genetic adaption of wastewater tolerance and stress adaption. The genome data and analyses presented in this study thus provides a fundamental resource for future studies on wastewater tolerance, secondary carotenoid accumulation, and algal stress biology, and will also pave a way for precise genome editing in target microalgal strains to engineeringly improve particular functionality in wastewater treatment.

## 3 Materials and Methods

### 3.1 Microalgal strains and culture system

Three industrial *Chlorella* genus strains were provided by Green Pac Bio Co (http://www.greenpacbio.com/), a biotechnology company aiming to develop new biological approaches for piggery wastewater treatment. *Chlorella sp.* Dachan were originally collected from the ditch sewage of ZhangHua city in Chinese Taipei and then cultivated in piggery wastewater. *Chlorella sorokiniana* BD09 and *Chlorella sorokiniana* BD08 were originated from *Chlorella sorokiniana* SU1, a lab collection at National Cheng Kung University and adapted to piggery wastewater. C*hlorella sorokiniana* UTEX 1602, also named FACHB-275, was purchased from FACHB-Collection, Institute of Hydrobiology, Chinese Academy of Sciences (Wuhan, China). Pure strain was obtained by serial transfers on solid BG11 Medium ^32^, maintained at 25°C, with 16: 8 day-dark external light illuminations at approximately 200 μmol m^−2^ s^−1^. The composition of BG11 medium was shown in Table S10.

### 3.2 Synthetic wastewater tolerance test

Four *Chlorella* strains, including BD08, BD09, Dachan, and UTEX 1602 underwent series of dilution to reach 100% purity and were then cultured in BG11 medium which contained 247 mg/L total nitrogen and 7.12 mg/L total phosphorus until the OD_680_ reach 5. Cells were then diluted into 50 ml synthetic wastewater at the same concentration as the initial stage (OD_680_=0.1). Three synthetic wastewater concentrations were used in the test: 500 mg/L glucose, 148.58 mg/L NH_4_Cl, 21.95 mg/L KH_2_PO_4_ on the basis of BG11 medium for high concentration treatment group; 300 mg/L glucose, 89.15 mg/L NH_4_Cl, 13.17 mg/L KH_2_PO_4_ on the basis of BG11 medium for middle concentration treatment group; 200 mg/L glucose, 59.43 mg/L NH_4_Cl, 8.78 mg/L KH_2_PO_4_ on the basis of BG11 medium for low concentration treatment group; and the pure BG11 medium was used as the control group. All treatments were performed in triplicate. The cultures were maintained at 25°C, with 16: 8 day-dark vs external light illumination cycle at approximately 200 μmol m^−2^ s^−1^. OD_680_ was measured each day to calculate growth rates of each strain. Bacterial antibiotics were added into the synthetic wastewater, for the purpose of avoiding bacterial contamination and disruption of OD_680_ measurement.

### 3.3 Genomic DNA extraction, library construction and genome sequencing

Microalgal cells were collected from liquid BG11 medium by centrifugation at 300 rpm at room temperature. Genomic DNA was then extracted using a modified cetyltrimethylammonium bromide (CTAB) method ^33^. The concentration of extracted DNA was determined by Qubit Fluorometer 2.0 (Life Technologies) and the quality was determined by electrophoresis analysis. A whole-genome shotgun strategy was applied to the qualified samples with large DNA fragments and adequate quantity, including pair-ended libraries with insert sizes ranging between 350 and 500 bp and mate-paired libraries with insert sizes of 2 kb, 5 kb, 10 kb and 20 kb, respectively. Whole genome sequencing was performed on the Illumina sequencers (HiSeq 2500 and HiSeq 4000) at BGI-Shenzhen.

To ensure high-quality of DNA sequencing reads (also called clean data) for downstream analyses, a strict filtering process was carried out using SOAPfilter (V2.2). Briefly, a series of filtering steps were performed on the raw reads, which include removal of Ns-rich (if Ns > 10%) reads, low quality reads (if > 10% of the bases for each read are defined as low-quality), reads with ≥ 10 bp aligned to the adapter sequences and PCR duplicates. Low quality read ends were also trimmed off by 5-8 bp.

### 3.4 RNA extraction, library construction and transcriptome sequencing

Algal cell mass was immediately frozen in liquid nitrogen after collection from liquid BG11 medium and stored at −80°C. Total RNA was isolated using a modified TRIZOL extraction protocol ^34^. The quantity and quality of RNA was determined by Agilent 2100 Bioanalyzer (Agilent Technologies).

For each RNA sample, mRNA was enriched using Dynabeads mRNA Purification Kit (Invitrogen), and then used to generate RNA-seq libraries after reverse transcription with BGI self-made reagents, with an insert size of ∼150 bp. Three RNA-seq libraries were barcoded and pooled together as the input to the BGISEQ-500 platform (BGI, Shenzhen) for sequencing. An average of 10 Gb clean data per sample was generated after filtering to ensure a complete set of expressed transcripts with sufficient coverage and depth for each sample.

### 3.5 Genome assembly and evaluation

Before the assembly, the HiSeq paired-end reads were used for the K-mer analysis to estimate the genome size ^13^. A K-mer is a sequence of length K from the clean reads. The peak of K-mer frequency (F) is determined by the total reads number (N), genome size (G), read length (L) and the length of K-mer (K) following the formula: F=N×(L-K+1)/G. The total K-mer number (M) is determined by the formula M=N×(L-K+1). As a result, the genome size was estimated as G=M/F. SOAPdenovo2 (version 2.04) ^35^ and Platanus (version 1.2.4) ^14^ were used for assembly. After several rounds of complete assemblies implementing alternative parameters, the optimal version of assembly was finalized by selecting the one with larger contig contiguity (estimated by contig N50) and higher genome completeness (estimated by RNA-seq mapping coverage) for downstream gap filling by Gapcloser (version 1.2) ^35^. Software MeDuSa (Multi-Draft based Scaffolder) ^36^ was used for genome scaffolding, which exploited information obtained from a set of draft genomes of three closely related species, *Chlorella variabilis* NC64A ^8^, *Chlorella sorokiniana* UTEX 1602 ^9^ and *Micractinium conductrix* SAG 241.80 ^9^. The completeness of the assembly was verified using BUSCO (benchmarking universal single-copy orthologues) analysis ^15^.

### 3.6 Gene prediction and valuation

For each genome assembly, repeat elements were masked before gene model prediction according to the instruction of RepeatMasker^37^. Gene models were predicted by MAKER-P pipeline (version 2.31) with two rounds of iterations ^38^. Briefly, the pipeline combined and integrated three methods. (1) Homolog-based method: Blast was used to align the genome assembly with an optimal core protein data set from four homologous species to assign the genes with putative functions. (2) *ab initio* prediction: a series of training was performed to obtain an optimal gene model prediction result. Augustus ^39^, SNAP ^40^, and Genemark-ES (version 4.21) ^41^ were used respectively for *ab initio* predictions. (3) Transcript-based method: EST sequences were obtained from transcriptome sequencing. Default parameters were used to run MAKER-P to combine and integrate all annotation sources, resulting in a high-quality set of gene models for each genome. The completeness of each set of predicted genes was evaluated using BUSCO analysis.

The functional annotation and assignment for each gene were performed against InterPro ^42^, KEGG, GO, COG, NR, SwissProt ^43^, and TrEMBL ^44^ databases.

### 3.7 Gene family clustering and phylogenetic analysis

Fourteen species were used for gene family clustering and phylogenetic construction in this study, including ten Chlorophyta (*Chlorella variabilis* NC64A ^8^, *Chlorella sorokiniana* UTEX 1602 ^9^, *Micractinium conductrix* SAG 241.80 ^9^, *Chlamydomonas reinhardtii* ^45^, *Micromonas commoda* ^46^, *Auxenochlorella protothecoides* ^10^, *Picochlorum sp.* SENEW3 ^47^, *Picochlorum costavermella* ^48^, *Ostreococcus tauri ^20^, Volvox carteri* ^49^), one Rhodophyta (*Cyanidioschyzon merolae* ^17^) and the three newly sequenced genomes in this study, BD09, BD08 and Dachan (see Table S9 for the version of the assembly). OrthoFinder ^16^ (inflation parameter: 1.3) was used to infer the orthogroups and orthologs based on the all-vs-all ncbi-blast-2.2.31+ results. Furthermore, we used the average amino acid bit score of the all-vs-all ncbi-blast-2.2.31+ results to generate a homologs similarity matrix to compare the identity of amino-acid sequences of these fourteen genomes.

Single-copy gene families (the family that every species only have one copy) extracted from the gene family clusters of the fourteen genomes were used to construct a phylogenetic tree. The alignments were performed by MAFFT ^50^ for each single-copy gene family, which were further concatenated into a super gene for each species. The aligned sites within the alignments were filtered out if >50% of the species were “gaps / Ns”. RAxML^51^ was then used to construct the maximum likelihood phylogenetic tree (with CAT+GTR amino acid substitution model, 500 bootstraps). The consensus phylogenetic species tree was plotted using FigTree v1.4.3 (http://tree.bio.ed.ac.uk/software/figtree/). All parameters were set with the default setting.

### 3.8 Comparative genomics within and between *Chlorella* genomes

Conserved syntenic and collinear blocks within and between genomes were detected by MCScanX ^18^. Scaffolds with larger than 1 Mb in length from all genomes were particularly selected for comparison across genomes in order to highlight and illustrate significant conserved genomic regions. Visualization of collinearity relationship was conducted using the Circos software ^19^. Synteny blocks were defined as the region where five or more conserved homologs were located. Syntenic region shared by two genomes were connected by colored lines.

### 3.9 Searching of candidate genes that were involved in nitrogen and phosphorus assimilation

Known candidate genes reported to be responsible for nitrogen and phosphorus assimilation were collected mainly from Emanuel Sanz-Luque ^24^ and Jeffrey Moseley ^52^. A strict criterion was set to define candidate genes. Firstly, Blastp (evalue <1e-10, similarity ≥ 80%, aligned coverage >80%) was used to search for candidate genes in target genomes using the known sequences as queries from *Chlamydomonas reinhardtii* or *Arabidopsis thaliana*. Further, only candidate genes that contain the functional assignment from SwissProt or NR database were retained as the final ones.

## 4 Conclusion

Our study provides three high-quality genome assemblies of industrial *Chlorella* genus with the highest assembly completeness and contiguity reported so far, compared with the other *Chlorella* clade genomes. We found candidate genes, gene families, and enriched metabolic pathways that are associated with the underlying molecular mechanism responsible for the fast-growing *Chlorella* strains adapted to wastewater. These data and analyses provide valuable information and insights into future work in genetic engineering and strain improvement of industrial wastewater-treatment *Chlorella*.

## Data Availability

All raw sequencing data of this study are available in the NCBI Sequence Read Archive (SRA) under accession number SRR8163051-SRR8163074. The whole genome assemblies for BD09, BD08 and Dachan in this study are deposited at DDBJ/ENA/GenBank under the accession numbers of RRZM00000000, RRZN00000000 and RRZL00000000, respectively. The data reported in this study are also available in the CNGB Nucleotide Sequence Archive (CNSA: http://db.cngb.org/cnsa; accession number CNP0000212).

## Acknowledgements

This work was funded by Science, Technology and Innovation Commission of Shenzhen Municipality under grant No. JCYJ20160531194327655 and No. JCYJ20151015162041454. This work was also supported by the Guangdong Provincial Key Laboratory of Genome Read and Write (No. 2017B030301011) and Guangdong Provincial Academician Workstation of BGI Synthetic Genomics (No. 2017B090904014).

## Author contributions

YG and YS conceived and designed the study. YY and YZS performed the experiment. TW, LZL, and XSJ analyzed the data.TW interpreted the data and drafted the manuscript. LC, XX and YG edited manuscript. All authors read and approved the final manuscript.

## Competing interests

The authors declare no competing interests.

## Notes

http://db.cngb.org/cnsa

