## Supplementary material for "Sequencing and comparative analysis of three *Chlorella* genomes provide insights into strain-specific adaptation to wastewater"

### 1. Supplementary Tables

**Table S1.** Actual piggery wastewater parameters and nutrient removal rate of *Chlorella sorokiniana* BD09 in actual piggery wastewater.

| Water quality parameters | Initial measurement | Measurement after 15 days | Nutrient removal rate of BD09 |
| --- | --- | --- | --- |
| COD (mg/L) | 2,652 | 731 | 72% |
| Total N (mg/L) | 321 | 39 | 88% |
| NH_4_-N (mg/L) | 271 | 38 | 86% |
| Total P (mg/L) | 44 | 17 | 61% |
| pH | 8.3 | 9.47 | / |

**Notes:**

1. Actual wastewater was diluted before *Chlorella* added.
2. Actual culture volume was 1.8 L, the initial OD_680_ of *Chlorella sorokiniana* BD09 was 1.

**Table S2.** Genomic and transcriptomic data statistics of three newly sequenced *Chlorella* strains.

| DNA/RNA data | Sample name | Insert size | Raw read length (bp) | Raw data (Gb) | Clean data (Gb) | Sequence depth of clean data (X) |
| --- | --- | --- | --- | --- | --- | --- |
| DNA | *Chlorella sorokiniana* BD09 | 350bp | 150_150 | 15.5 | 10.56 | 195.6 |
|  |  | 500bp | 250_250 | 25.3 | 17.28 | 320 |
|  |  | 2kb | 100_100 | 12.4 | 3.79 | 70.2 |
|  |  | 5kb | 100_100 | 9.8 | 5.31 | 98.3 |
|  |  | 10Kb | 100_100 | 6.3 | 3.56 | 65.9 |
|  |  | 20kb | 100_100 | 6.4 | 2.66 | 49.3 |
|  | Total | / | / | 75.7 | 43.16 | 799.3 |
|  | *Chlorella sorokiniana* BD08 | 350bp | 150_150 | 15.5 | 9.49 | 161.9 |
|  |  | 500bp | 250_250 | 25.3 | 16.34 | 278.8 |
|  |  | 2kb | 100_100 | 12.4 | 6.66 | 113.7 |
|  |  | 5kb | 100_100 | 9.8 | 5.15 | 87.9 |
|  |  | 10Kb | 100_100 | 6.3 | 3.59 | 61.3 |
|  |  | 20kb | 100_100 | 6.4 | 2.75 | 46.9 |
|  | Total | / | / | 75.7 | 43.98 | 750.5 |
|  | *Chlorella sp.* Dachan | 350bp | 150_150 | 14.6 | 9.72 | 160.9 |
|  |  | 500bp | 250_250 | 23.5 | 15.59 | 258.1 |
|  |  | 2kb | 100_100 | 10.7 | 4.06 | 67.2 |
|  |  | 5kb | 100_100 | 10.3 | 5.72 | 94.7 |
|  |  | 10Kb | 100_100 | 5.7 | 3.12 | 51.7 |
|  |  | 20kb | 100_100 | 6.6 | 2.88 | 47.7 |
|  | Total | / | / | 71.4 | 41.09 | 680.3 |
| RNA | *Chlorella sorokiniana* BD09 | 150bp | 100_100 | 23.5 | 14.4 | / |
|  | *Chlorella sorokiniana* BD08 | 150bp | 100_100 | 19.1 | 11.6 | / |
|  | *Chlorella sp.* Dachan | 150bp | 100_100 | 22.9 | 12.6 | / |

**Table S3.** A summary of scaffolds length distribution of our three newly sequenced genomes *Chlorella sorokiniana* BD09, *Chlorella sorokiniana* BD08 and *Chlorella sp.* Dachan in comparison with other three published *Chlorella* clade genomes.

|  | *Chlorella sorokiniana*  BD08 | | *Chlorella sorokiniana* BD09 | | *Chlorella sp.* Dachan | | *Chlorella variabilis*  NC64A | | *Chlorella sorokiniana* UTEX 1602 | | Micractinium *conductrix* | |
| --- | --- | --- | --- | --- | --- | --- | --- | --- | --- | --- | --- | --- |
| **Scaffold length (bp)** | **No.** | **Size (bp)** | **No.** | **Size (bp)** | **No.** | **Size (bp)** | **No.** | **Size (bp)** | **No.** | **Size (bp)** | **No.** | **Size (bp)** |
| ≥ 10,000 | 33 | 57,379,143 | 20 | 53,438,449 | 46 | 58,259,782 | 82 | 45,222,670 | 139 | 59,422,181 | 246 | 60,680,316 |
| ≥ 8,000 | 38 | 57,424,705 | 22 | 53,457,128 | 55 | 58,344,829 | 95 | 45,338,289 | 148 | 59,505,254 | 262 | 60,823,285 |
| ≥ 5,000 | 43 | 57,455,370 | 23 | 53,463,667 | 72 | 58,454,003 | 134 | 45,602,803 | 155 | 59,552,060 | 283 | 60,962,532 |
| ≥ 4,000 | 52 | 57,494,367 | 26 | 53,477,742 | 90 | 58,535,294 | 153 | 45,688,397 | 156 | 59,556,856 | 288 | 60,985,293 |
| ≥ 2,000 | 137 | 57,734,923 | 45 | 53,527,384 | 214 | 58,893,416 | 235 | 45,930,001 | 158 | 59,564,266 | 298 | 61,015,730 |
| ≥ 1,000 | 537 | 58,306,158 | 93 | 53,589,812 | 711 | 59,594,959 | 414 | 46,159,512 | 159 | 59,566,223 | 300 | 61,018,900 |
| ≥ 800 | 671 | 58,425,778 | 130 | 53,623,002 | 909 | 59,771,270 | / | / | / | / | / | / |
| ≥ 600 | 819 | 58,529,408 | 325 | 53,757,022 | 1,227 | 59,994,888 | / | / | / | / | / | / |
| ≥ 500 | 905 | 58,576,441 | 427 | 53,813,149 | 1,385 | 60,082,138 | / | / | / | / | / | / |
| ≥ 300 | 1,104 | 58,653,236 | 679 | 53,912,376 | 1,852 | 60,263,378 | / | / | / | / | / | / |
| ≥ 200 | 1,242 | 58,686,758 | 920 | 53,971,092 | 2,277 | 60,366,751 | / | / | / | / | / | / |
| **Total** | 1,242 | 58,686,758 | 920 | 53,971,092 | 2,277 | 60,366,751 | 414 | 46,159,512 | 159 | 59,566,223 | 300 | 61,018,900 |

**Table S4.** The proportion of different types of repeat sequences (%) in the three newly sequenced *Chlorella* strains.

| Repeat elements | *Chlorella sorokiniana* BD09 | | *Chlorella sorokiniana* BD08 | | *Chlorella sp.* Dachan | |
| --- | --- | --- | --- | --- | --- | --- |
|  | Repeat size (bp) | % of genome | Repeat size (bp) | % of genome | Repeat size (bp) | % of genome |
| LTR | 747,570 | 1.37 | 813,108 | 1.38 | 1,361,472 | 2.12 |
| LINEs | 710,610 | 1.31 | 734,058 | 1.24 | 770,850 | 1.2 |
| SINEs | 22,063 | 0.04 | 4,454 | 0 | 16,088 | 0.02 |
| Total DNA-TEs | 111,042 | 0.2 | 210,928 | 0.35 | 198,442 | 0.3 |
| Satellite | 1,388 | 0 | 754 | 0 | 1,993 | 0 |
| Simple repeats | 43,684 | 0.08 | 46,758 | 0.07 | 88,870 | 0.13 |
| Other | 1,970 | 0 | 1,912 | 0 | 1,217 | 0 |
| Unknown | 499,375 | 0.92 | 517,631 | 0.88 | 838,393 | 1.3 |
| **Total** | 1927537 | 3.56 | 2136707 | 3.63 | 3047886 | 4.76 |

**Table S5.** Number of genes with functional annotation against seven different databases in three newly sequenced *Chlorella* strains.

|  |  | *Chlorella sorokiniana* BD09 | | *Chlorella sorokiniana* BD08 | | *Chlorella sp.* Dachan Dachan | |
| --- | --- | --- | --- | --- | --- | --- | --- |
| Type | Database | Gene number | Percentage (%) | Gene number | Percentage (%) | Gene number | Percentage (%) |
| Total genes | / | 9,668 | 100% | 10,240 | 100% | 9,821 | 100% |
| Annotated | Nr | 8,550 | 88.44% | 8,909 | 86.99% | 8,483 | 86.38% |
|  | Swissprot | 6,107 | 63.17% | 6,229 | 60.82% | 5,934 | 60.42% |
|  | KEGG | 5,943 | 61.47% | 6,112 | 59.68% | 5,819 | 59.25% |
|  | COG | 4,427 | 45.79% | 4,485 | 43.79% | 4,318 | 43.97% |
|  | TrEMBL | 8,587 | 88.82% | 8.960 | 87.49% | 8,501 | 86.56% |
|  | Interpro | 7,139 | 73.84% | 7,346 | 71.73% | 7,012 | 71.40% |
|  | GO | 4,039 | 41.78% | 4,137 | 40.40% | 3,999 | 40.72% |
| All annotated | / | 8,720 | 90.19% | 9,128 | 89.13% | 8,678 | 88.36% |

**Table S6**. KEGG pathway enrichment of genes that were expanded in *Chlorella sorokiniana* BD09 compared to *Chlorella variabilis* NC64A (p-value < 0.05).

| # | Pathway | DEGs with pathway annotation (612) | All genes with pathway annotation (5943) | P-value | Q-value | Pathway ID |
| --- | --- | --- | --- | --- | --- | --- |
| 1 | Endocytosis | 25 (4.08%) | 117 (1.97%) | 0.0002763611 | 0.02708339 | ko04144 |
| 2 | Plant hormone signal transduction | 27 (4.41%) | 138 (2.32%) | 0.0007063922 | 0.03461322 | ko04075 |
| 3 | Starch and sucrose metabolism | 17 (2.78%) | 83 (1.4%) | 0.004053814 | 0.13242459 | ko00500 |
| 4 | Stilbenoid, diarylheptanoid and gingerol biosynthesis | 3 (0.49%) | 6 (0.1%) | 0.01712779 | 0.41963086 | ko00945 |
| 5 | Other glycan degradation | 4 (0.65%) | 12 (0.2%) | 0.02811379 | 0.55103028 | ko00511 |

**Table S7.** KEGG pathway enrichment of genes that were shared among *Chlorella sorokiniana* BD09, *Chlorella sorokiniana* BD08 and *Chlorella sp.* Dachan compared to *Chlorella variabilis* NC64A (p-value < 0.05).

| # | Pathway | DEGs with pathway annotation (2085) | All genes with pathway annotation (17874) | P-value | Q-value | Pathway ID |
| --- | --- | --- | --- | --- | --- | --- |
| 1 | Plant hormone signal transduction | 85 (4.08%) | 378 (2.11%) | 1.3985e-09 | 1.510380e-07 | ko04075 |
| 2 | Endocytosis | 62 (2.97%) | 337 (1.89%) | 0.0001758213 | 6.768943e-03 | ko04144 |
| 3 | Fatty acid elongation | 15 (0.72%) | 47 (0.26%) | 0.0001880262 | 6.768943e-03 | ko00062 |
| 4 | Stilbenoid, diarylheptanoid and gingerol biosynthesis | 7 (0.34%) | 14 (0.08%) | 0.0004736567 | 1.278873e-02 | ko00945 |
| 5 | Nitrogen metabolism | 21 (1.01%) | 99 (0.55%) | 0.004563128 | 9.856356e-02 | ko00910 |
| 6 | Isoflavonoid biosynthesis | 3 (0.14%) | 4 (0.02%) | 0.005786986 | 1.041657e-01 | ko00943 |
| 7 | Other glycan degradation | 9 (0.43%) | 31 (0.17%) | 0.007233218 | 1.115982e-01 | ko00511 |
| 8 | Diterpenoid biosynthesis | 5 (0.24%) | 13 (0.07%) | 0.0123529 | 1.586051e-01 | ko00904 |
| 9 | Arachidonic acid metabolism | 16 (0.77%) | 76 (0.43%) | 0.01321709 | 1.586051e-01 | ko00590 |
| 10 | Phosphonate and phosphinate metabolism | 4 (0.19%) | 10 (0.06%) | 0.02171426 | 1.956423e-01 | ko00440 |
| 11 | Cutin, suberine and wax biosynthesis | 4 (0.19%) | 10 (0.06%) | 0.02171426 | 1.956423e-01 | ko00073 |
| 12 | Starch and sucrose metabolism | 38 (1.82%) | 234 (1.31%) | 0.02173803 | 1.956423e-01 | ko00500 |
| 13 | Ether lipid metabolism | 11 (0.53%) | 51 (0.29%) | 0.03075869 | 2.234067e-01 | ko00565 |
| 14 | Zeatin biosynthesis | 4 (0.19%) | 11 (0.06%) | 0.03102871 | 2.234067e-01 | ko00908 |
| 15 | Lipoic acid metabolism | 4 (0.19%) | 11 (0.06%) | 0.03102871 | 2.234067e-01 | ko00785 |

**Table S8.** KEGG pathway enrichment of genes in the inversion region of *Chlorella sorokiniana* BD09, *Chlorella sorokiniana* BD08 and *Chlorella sp.* Dachan genomes (p-value < 0.05).

| Strains | # | Pathway | DEGs with pathway annotation (202) | All genes with pathway annotation (5943) | P-value | Q-value | Pathway ID |
| --- | --- | --- | --- | --- | --- | --- | --- |
| BD09 | 1 | DNA replication | 7 (3.47%) | 59 (0.99%) | 0.003596674 | 0.2553639 | ko03030 |
|  | 2 | Phenylalanine, tyrosine and tryptophan biosynthesis | 4 (1.98%) | 29 (0.49%) | 0.01580532 | 0.4477446 | ko00400 |
|  | 3 | Protein export | 4 (1.98%) | 33 (0.56%) | 0.02451788 | 0.4477446 | ko03060 |
|  | 4 | RNA transport | 10 (4.95%) | 151 (2.54%) | 0.03232945 | 0.4477446 | ko03013 |
|  | 5 | RNA polymerase | 4 (1.98%) | 36 (0.61%) | 0.03263405 | 0.4477446 | ko03020 |
|  | 6 | Pyrimidine metabolism | 8 (3.96%) | 113 (1.9%) | 0.03783757 | 0.4477446 | ko00240 |
|  | 7 | mRNA surveillance pathway | 6 (2.97%) | 77 (1.3%) | 0.04611195 | 0.4677069 | ko03015 |
| BD08 | 1 | Phenylalanine, tyrosine and tryptophan biosynthesis | 6 (1.66%) | 25 (0.41%) | 0.002800962 | 0.1311842 | ko00400 |
|  | 2 | DNA replication | 10 (2.76%) | 65 (1.06%) | 0.004509143 | 0.1311842 | ko03030 |
|  | 3 | Plant hormone signal transduction | 17 (4.7%) | 144 (2.36%) | 0.004741597 | 0.1311842 | ko04075 |
|  | 4 | Protein export | 5 (1.38%) | 29 (0.47%) | 0.02595792 | 0.5386268 | ko03060 |
|  | 5 | Other types of O-glycan biosynthesis | 5 (1.38%) | 33 (0.54%) | 0.04282485 | 0.7108925 | ko00514 |
| Dachan | 1 | DNA replication | 7 (4.05%) | 64 (1.1%) | 0.002707272 | 0.1597290 | ko03030 |
|  | 2 | Protein export | 4 (2.31%) | 30 (0.52%) | 0.01130525 | 0.3335049 | ko03060 |
|  | 3 | Spliceosome | 9 (5.2%) | 143 (2.46%) | 0.02605555 | 0.4425155 | ko03040 |
|  | 4 | Base excision repair | 4 (2.31%) | 40 (0.69%) | 0.03000105 | 0.4425155 | ko03410 |
|  | 5 | Pyrimidine metabolism | 7 (4.05%) | 110 (1.89%) | 0.04473997 | 0.4518410 | ko00240 |
|  | 6 | Homologous recombination | 4 (2.31%) | 47 (0.81%) | 0.0499743 | 0.4518410 | ko03440 |

**Table S9.** Species list and genome versions used for comparative genomics analysis.

| Species | Data Sources |
| --- | --- |
| *Auxenochlorella protothecoides* | NCBI (ASM73321v1) |
| *Cyanidioschyzon merolae* | NCBI (ASM9120v1) |
| *Micromonas commoda* | Ghent University |
| *Chlamydomonas reinhardtii* | [DOE Joint Genome Institute](http://genome.jgi-psf.org/Chlre3/Chlre3.home.html) [( GCA_000002595.3)](https://www.ncbi.nlm.nih.gov/genome/147?genome_assembly_id=22881) |
| *Volvox carteri* | DOE Joint Genome Institute  [(GCA_000143455.1)](https://www.ncbi.nlm.nih.gov/genome/413?genome_assembly_id=29051) |
| *Ostreococcus tauri* | Ghent University |
| *Picochlorum sp.* SENEW3 | NCBI (ASM8764v1) |
| *Picochlorum costavermella* | ORCAE (20151222) |
| *Chlorella variabilis* NC64A | DOE Joint Genome Institute (GCA_000147415.1) |
| *Chlorella sorokiniana* UTEX 1602 | NCBI (ASM313072v1) |
| *Micractinium conductrix* SAG 241.80 | NCBI (ASM224581v2) |

**Table S10.** The composition of BG11 medium.

| **Compound** | **Concentration (mg/liter)** |
| --- | --- |
| NaNO_3_ | 1,500 |
| K_2_HPO_4_ | 40 |
| MgSO_4_.7H_2_O | 75 |
| CaCl_2_.2H_2_O | 36 |
| Citric acid | 6 |
| Ammonium ferric citrate green | 6 |
| EDTANa_2_ | 1 |
| Na_2_CO_3_ | 20 |
| Trace metal solution A5 | 1 ml/liter |

The detailed information of A5:

| **Compound** | **Concentration (mg/liter)** |
| --- | --- |
| H_3_BO_3_ | 2,860 |
| MnCl_2_.4H_2_O | 1,810 |
| ZnSO_4_.7H_2_O | 222 |
| Na_2_MoO_4_.2H_2_O | 390 |
| CuSO_4_.5H_2_O | 79 |
| Co(NO_3_)_2_.6H_2_O | 494 |

Adjust pH to 7.1 with 1M NaOH or HCl.

### 2. Supplementary Figures

| **a** |
| --- |
| **b** |
| **c** |

Chlorella sorokiniana BD09

| 17-mer depth | Frequency (%) |
| --- | --- |
| 1 | 39.0379 |
| 2 | 1.73845 |
| 3 | 0.313828 |
| 4 | 0.17655 |
| 5 | 0.143437 |
| 6 | 0.133323 |
| 7 | 0.131264 |
| 8 | 0.134339 |
| 9 | 0.141901 |
| 10 | 0.146314 |
| 11 | 0.152468 |
| 12 | 0.162389 |
| 13 | 0.175955 |
| 14 | 0.186055 |
| 15 | 0.197915 |
| 16 | 0.213642 |
| 17 | 0.227112 |
| 18 | 0.241588 |
| 19 | 0.259558 |
| 20 | 0.277013 |
| 21 | 0.29628 |
| 22 | 0.317272 |
| 23 | 0.337913 |
| 24 | 0.358458 |
| 25 | 0.381612 |
| 26 | 0.408704 |
| 27 | 0.440928 |
| 28 | 0.480256 |
| 29 | 0.536495 |
| 30 | 0.600801 |
| 31 | 0.678478 |
| 32 | 0.77064 |
| 33 | 0.883815 |
| 34 | 1.00616 |
| 35 | 1.14552 |
| 36 | 1.30044 |
| 37 | 1.46633 |
| 38 | 1.63688 |
| 39 | 1.79963 |
| 40 | 1.95136 |
| 41 | 2.09177 |
| 42 | 2.20592 |
| 43 | 2.293 |
| 44 | 2.34405 |
| 45 | 2.36321 |
| 46 | 2.34634 |
| 47 | 2.29582 |
| 48 | 2.20899 |
| 49 | 2.09756 |
| 50 | 1.9563 |
| 51 | 1.80096 |
| 52 | 1.63385 |
| 53 | 1.46187 |
| 54 | 1.28997 |
| 55 | 1.12446 |
| 56 | 0.967794 |
| 57 | 0.822757 |
| 58 | 0.69254 |
| 59 | 0.577116 |
| 60 | 0.476201 |
| 61 | 0.390812 |
| 62 | 0.319004 |
| 63 | 0.259225 |
| 64 | 0.209393 |
| 65 | 0.169733 |
| 66 | 0.138666 |
| 67 | 0.11451 |
| 68 | 0.097342 |
| 69 | 0.0835076 |
| 70 | 0.0732653 |
| 71 | 0.0674797 |
| 72 | 0.0633045 |
| 73 | 0.0601209 |
| 74 | 0.0592353 |
| 75 | 0.0592913 |
| 76 | 0.0604422 |
| 77 | 0.0628599 |
| 78 | 0.0645563 |
| 79 | 0.067111 |
| 80 | 0.0695323 |
| 81 | 0.0729975 |
| 82 | 0.0759445 |
| 83 | 0.0785216 |
| 84 | 0.081146 |
| 85 | 0.0834739 |
| 86 | 0.0848453 |
| 87 | 0.0864832 |
| 88 | 0.0879243 |
| 89 | 0.0886941 |
| 90 | 0.0890852 |
| 91 | 0.0893181 |
| 92 | 0.0886542 |
| 93 | 0.0871894 |
| 94 | 0.0854643 |
| 95 | 0.0837579 |
| 96 | 0.0811397 |
| 97 | 0.0774242 |
| 98 | 0.074552 |
| 99 | 0.0708402 |
| 100 | 0.0663325 |
| 101 | 0.0631912 |
| 102 | 0.0593698 |
| 103 | 0.0549979 |
| 104 | 0.0518404 |
| 105 | 0.0473439 |
| 106 | 0.043941 |
| 107 | 0.040436 |
| 108 | 0.0367541 |
| 109 | 0.0336128 |
| 110 | 0.0313359 |
| 111 | 0.0286891 |
| 112 | 0.0262403 |
| 113 | 0.0241452 |
| 114 | 0.0221162 |
| 115 | 0.0204173 |
| 116 | 0.0191505 |
| 117 | 0.018027 |
| 118 | 0.0169932 |
| 119 | 0.016343 |
| 120 | 0.0155272 |
| 121 | 0.0151174 |
| 122 | 0.0147686 |
| 123 | 0.014603 |
| 124 | 0.0141284 |
| 125 | 0.0140375 |
| 126 | 0.0140861 |
| 127 | 0.014258 |
| 128 | 0.0143327 |
| 129 | 0.0141695 |
| 130 | 0.0139839 |
| 131 | 0.0141857 |
| 132 | 0.0140512 |
| 133 | 0.0141446 |
| 134 | 0.0141309 |
| 135 | 0.0143115 |
| 136 | 0.0140624 |
| 137 | 0.0139092 |
| 138 | 0.0139702 |
| 139 | 0.0138793 |
| 140 | 0.0135679 |
| 141 | 0.0132266 |
| 142 | 0.013051 |
| 143 | 0.012985 |
| 144 | 0.0127122 |
| 145 | 0.0121181 |
| 146 | 0.0120471 |
| 147 | 0.0116498 |
| 148 | 0.0111428 |
| 149 | 0.0107866 |
| 150 | 0.0103656 |

Chlorella sorokiniana BD08

| 17-mer depth | Frequency (%) |
| --- | --- |
| 1 | 49.4394 |
| 2 | 3.04411 |
| 3 | 0.623464 |
| 4 | 0.324894 |
| 5 | 0.246129 |
| 6 | 0.228309 |
| 7 | 0.227635 |
| 8 | 0.238693 |
| 9 | 0.260063 |
| 10 | 0.284051 |
| 11 | 0.315465 |
| 12 | 0.350898 |
| 13 | 0.396278 |
| 14 | 0.45345 |
| 15 | 0.5331 |
| 16 | 0.637915 |
| 17 | 0.775148 |
| 18 | 0.952897 |
| 19 | 1.17141 |
| 20 | 1.41345 |
| 21 | 1.68349 |
| 22 | 1.9536 |
| 23 | 2.20553 |
| 24 | 2.43035 |
| 25 | 2.59322 |
| 26 | 2.6813 |
| 27 | 2.69689 |
| 28 | 2.63852 |
| 29 | 2.50016 |
| 30 | 2.30427 |
| 31 | 2.07154 |
| 32 | 1.81503 |
| 33 | 1.55056 |
| 34 | 1.29544 |
| 35 | 1.05692 |
| 36 | 0.842178 |
| 37 | 0.661023 |
| 38 | 0.511242 |
| 39 | 0.390391 |
| 40 | 0.297411 |
| 41 | 0.230914 |
| 42 | 0.18182 |
| 43 | 0.147862 |
| 44 | 0.125877 |
| 45 | 0.111321 |
| 46 | 0.10424 |
| 47 | 0.101453 |
| 48 | 0.101434 |
| 49 | 0.10337 |
| 50 | 0.103999 |
| 51 | 0.106065 |
| 52 | 0.107618 |
| 53 | 0.108641 |
| 54 | 0.108607 |
| 55 | 0.107688 |
| 56 | 0.105138 |
| 57 | 0.101999 |
| 58 | 0.0971757 |
| 59 | 0.0922289 |
| 60 | 0.0857224 |
| 61 | 0.0797017 |
| 62 | 0.0731598 |
| 63 | 0.0660741 |
| 64 | 0.0601596 |
| 65 | 0.05378 |
| 66 | 0.0482766 |
| 67 | 0.0430427 |
| 68 | 0.0388935 |
| 69 | 0.0351259 |
| 70 | 0.031908 |
| 71 | 0.0291426 |
| 72 | 0.0267036 |
| 73 | 0.0249698 |
| 74 | 0.0233982 |
| 75 | 0.0220548 |
| 76 | 0.0211353 |
| 77 | 0.0206043 |
| 78 | 0.019968 |
| 79 | 0.0195048 |
| 80 | 0.0191517 |
| 81 | 0.0185439 |
| 82 | 0.0183256 |
| 83 | 0.0175703 |
| 84 | 0.0172576 |
| 85 | 0.0168426 |
| 86 | 0.0162908 |
| 87 | 0.0157372 |
| 88 | 0.0154874 |
| 89 | 0.0148137 |
| 90 | 0.0141233 |
| 91 | 0.0133523 |
| 92 | 0.0123984 |
| 93 | 0.0118496 |
| 94 | 0.0111405 |
| 95 | 0.0105849 |
| 96 | 0.00984138 |
| 97 | 0.00926115 |
| 98 | 0.00900348 |
| 99 | 0.00850389 |
| 100 | 0.008141 |
| 101 | 0.00763649 |
| 102 | 0.00722737 |
| 103 | 0.00709854 |
| 104 | 0.00681236 |
| 105 | 0.0064534 |
| 106 | 0.00618983 |
| 107 | 0.00602461 |

Chlorella sp. Dachan

| 17-mer depth | Frequency (%) |
| --- | --- |
| 1 | 3.266 |
| 2 | 1.58618 |
| 3 | 0.399413 |
| 4 | 0.246191 |
| 5 | 0.204845 |
| 6 | 0.191225 |
| 7 | 0.191324 |
| 8 | 0.192564 |
| 9 | 0.19648 |
| 10 | 0.20315 |
| 11 | 0.211439 |
| 12 | 0.224718 |
| 13 | 0.239003 |
| 14 | 0.253486 |
| 15 | 0.273195 |
| 16 | 0.295409 |
| 17 | 0.318984 |
| 18 | 0.342906 |
| 19 | 0.368581 |
| 20 | 0.394234 |
| 21 | 0.42298 |
| 22 | 0.45186 |
| 23 | 0.480995 |
| 24 | 0.517698 |
| 25 | 0.554505 |
| 26 | 0.595163 |
| 27 | 0.647783 |
| 28 | 0.706767 |
| 29 | 0.778259 |
| 30 | 0.863667 |
| 31 | 0.962657 |
| 32 | 1.08375 |
| 33 | 1.22032 |
| 34 | 1.36342 |
| 35 | 1.51755 |
| 36 | 1.68015 |
| 37 | 1.8438 |
| 38 | 2.00576 |
| 39 | 2.14983 |
| 40 | 2.27789 |
| 41 | 2.37903 |
| 42 | 2.44824 |
| 43 | 2.48882 |
| 44 | 2.49173 |
| 45 | 2.46373 |
| 46 | 2.40142 |
| 47 | 2.29568 |
| 48 | 2.17212 |
| 49 | 2.02336 |
| 50 | 1.85046 |
| 51 | 1.67577 |
| 52 | 1.49399 |
| 53 | 1.31243 |
| 54 | 1.13945 |
| 55 | 0.981734 |
| 56 | 0.82981 |
| 57 | 0.695481 |
| 58 | 0.579287 |
| 59 | 0.47498 |
| 60 | 0.388165 |
| 61 | 0.315992 |
| 62 | 0.256956 |
| 63 | 0.210068 |
| 64 | 0.172108 |
| 65 | 0.142516 |
| 66 | 0.120574 |
| 67 | 0.103251 |
| 68 | 0.0907293 |
| 69 | 0.0826425 |
| 70 | 0.0762133 |
| 71 | 0.0731187 |
| 72 | 0.0721961 |
| 73 | 0.0709005 |
| 74 | 0.0718987 |
| 75 | 0.07341 |
| 76 | 0.0757794 |
| 77 | 0.0781829 |
| 78 | 0.0801976 |
| 79 | 0.0827583 |
| 80 | 0.0854299 |
| 81 | 0.0873678 |
| 82 | 0.0888048 |
| 83 | 0.0915349 |
| 84 | 0.0928878 |
| 85 | 0.0945503 |
| 86 | 0.0957032 |
| 87 | 0.0954717 |
| 88 | 0.0957727 |
| 89 | 0.094532 |
| 90 | 0.094109 |
| 91 | 0.0917848 |
| 92 | 0.0891168 |
| 93 | 0.0864781 |
| 94 | 0.0840185 |
| 95 | 0.0817869 |
| 96 | 0.0778855 |
| 97 | 0.0738208 |
| 98 | 0.0699279 |
| 99 | 0.0666213 |
| 100 | 0.0622774 |
| 101 | 0.0575533 |
| 102 | 0.0536458 |
| 103 | 0.0490802 |
| 104 | 0.0457236 |
| 105 | 0.0426766 |
| 106 | 0.0386252 |
| 107 | 0.0354661 |
| 108 | 0.0324764 |
| 109 | 0.0306481 |
| 110 | 0.02784 |
| 111 | 0.0258863 |
| 112 | 0.0238947 |
| 113 | 0.0220251 |
| 114 | 0.0207636 |
| 115 | 0.0192291 |
| 116 | 0.0183918 |
| 117 | 0.0177605 |
| 118 | 0.0174643 |
| 119 | 0.0166684 |
| 120 | 0.0164758 |
| 121 | 0.0159237 |
| 122 | 0.0157031 |
| 123 | 0.0153777 |
| 124 | 0.0151632 |
| 125 | 0.0149938 |
| 126 | 0.0150901 |
| 127 | 0.015162 |
| 128 | 0.015162 |
| 129 | 0.0149133 |
| 130 | 0.0151815 |
| 131 | 0.014817 |
| 132 | 0.0149097 |
| 133 | 0.014644 |
| 134 | 0.014672 |
| 135 | 0.0143783 |
| 136 | 0.0143892 |
| 137 | 0.0142418 |
| 138 | 0.0141955 |
| 139 | 0.0135763 |
| 140 | 0.0133496 |
| 141 | 0.0129145 |
| 142 | 0.0125354 |
| 143 | 0.0123977 |
| 144 | 0.0117432 |
| 145 | 0.0113703 |
| 146 | 0.0109754 |
| 147 | 0.0107889 |
| 148 | 0.010249 |
| 149 | 0.010043 |
| 150 | 0.00942504 |
| 151 | 0.00950061 |
| 152 | 0.0088193 |
| 153 | 0.00856335 |
| 154 | 0.00827327 |
| 155 | 0.00790641 |
| 156 | 0.00758221 |
| 157 | 0.0075042 |
| 158 | 0.00729335 |
| 159 | 0.00707274 |
| 160 | 0.00701424 |
| 161 | 0.00686676 |
| 162 | 0.00654378 |
| 163 | 0.00630855 |
| 164 | 0.00610257 |
| 165 | 0.00581615 |
| 166 | 0.00596728 |
| 167 | 0.0057418 |
| 168 | 0.00553826 |
| 169 | 0.00559555 |
| 170 | 0.00565649 |
| 171 | 0.0053981 |
| 172 | 0.00544198 |
| 173 | 0.00555533 |
| 174 | 0.00532253 |
| 175 | 0.0053006 |
| 176 | 0.00508365 |
| 177 | 0.00505805 |
| 178 | 0.00488864 |
| 179 | 0.00493252 |
| 180 | 0.00501174 |
| 181 | 0.00484354 |
| 182 | 0.00486426 |
| 183 | 0.00471435 |
| 184 | 0.00465463 |
| 185 | 0.00446693 |
| 186 | 0.00443037 |
| 187 | 0.00444865 |
| 188 | 0.00446449 |
| 189 | 0.00428899 |
| 190 | 0.00417686 |
| 191 | 0.00405619 |
| 192 | 0.00407204 |
| 193 | 0.00414882 |
| 194 | 0.00396235 |
| 195 | 0.00399891 |
| 196 | 0.00385997 |
| 197 | 0.0037198 |
| 198 | 0.0036613 |
| 199 | 0.00362961 |
| 200 | 0.00352723 |

**Figure S1**. 17-kmer estimation of genome sizes among three newly sequenced *Chlorella* strains. The x-axis was depth (X); the y-axis was K-mer frequency. a) The genome size of *Chlorella sorokiniana* BD09 was estimated to be 55.1 Mb based on reads from short insert size library. b) The genome size of *Chlorella sorokiniana* BD08 was estimated to be 56.5 Mb based on reads from short insert size library. c) The genome size of *Chlorella sp.* Dachan was estimated to be 60.4 Mb based on reads from short insert size library.

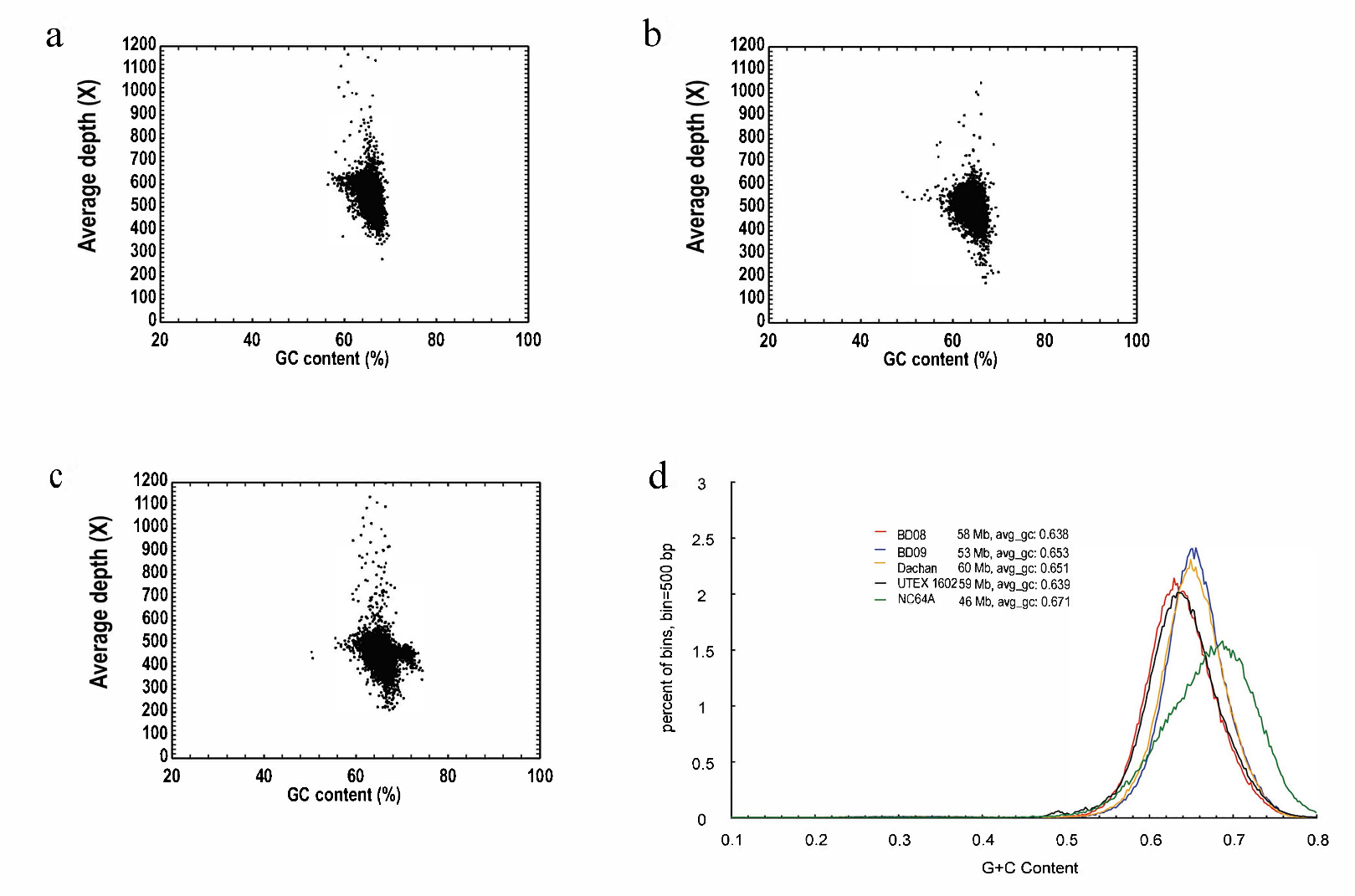

**Figure S2.** GC content analysis of three newly sequenced genomes (a, b, c) and comparison of the GC contents across five *Chlorella* genomes (d). a) GC content analysis of *Chlorella sorokiniana* BD09; b) GC content analysis of *Chlorella sorokiniana* BD08; c) GC content analysis of *Chlorella sp.* Dachan. In the figure d, the x-axis was GC content and the y-axis indicated the proportion of the bin number divided by the total windows. We used 500 bp bins (with 250 bp overlap) sliding along the genome. The GC content of three genomes was 65.3% (BD09), 63.8% (BD08) and 65.1% (Dachan) respectively, which was similar to *Chlorella sorokiniana* UETX 1602 (63.9%). All four genomes were GC rich that slightly lower than *Chlorella variabilis* NC64A (67.1%).

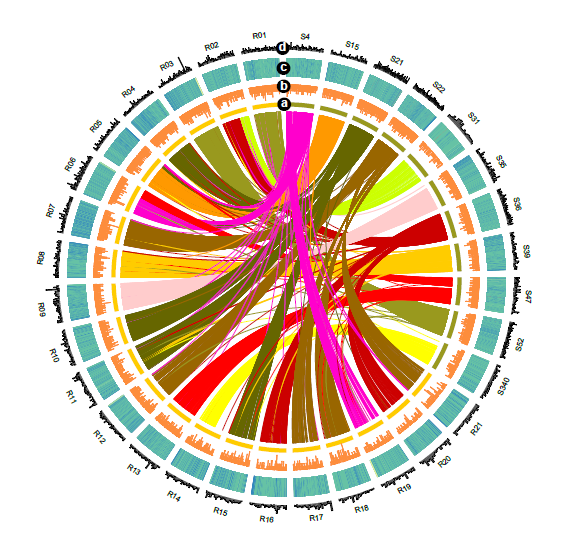

**Figure S3.** Circos visualization of collinearity between *Chlorella sorokiniana* BD08 (S) and *Chlorella sorokiniana* UTEX 1602 (R), only scaffolds with length >1Mb were used. Different layers denoted: 1) collinear regions between the two strains connected by colored lines, 2) gene density, 3) GC content, and 4) TE density.

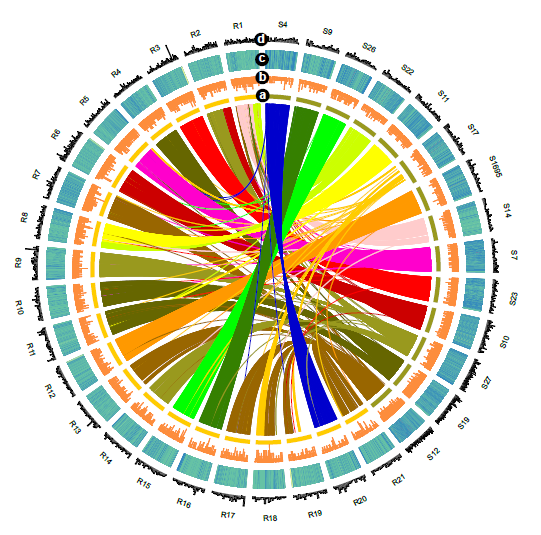

**Figure S4.** Circos visualization of collinearity between *Chlorella sorokiniana* BD09 (S) and *Chlorella sorokiniana* UTEX 1602 (R), only scaffolds with length >1Mb were used. Different layers denoted: 1) collinear regions between the two strains connected by colored lines, 2) gene density, 3) GC content, and 4) TE density.

| a | 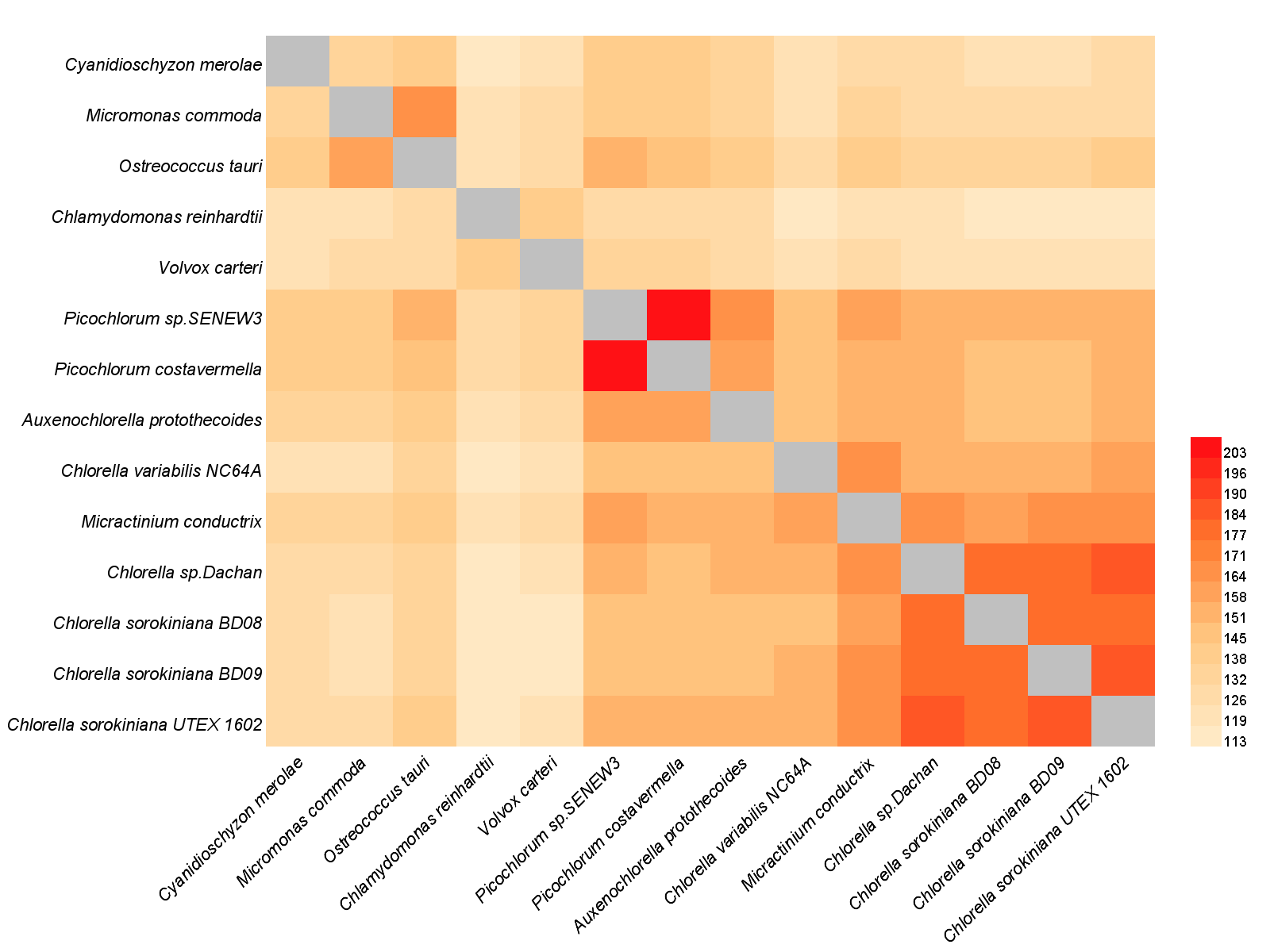 |
| --- | --- |
| b |  |

Identity of mino-acid sequences

|  | Similartity (%) |
| --- | --- |
| Cyanidioschyzon merolae | 50.9883% |
| Micromonas commoda | 50.2239% |
| Ostreococcus tauri | 54.0959% |
| Chlamydomonas reinhardtii | 45.2932% |
| Volvox carteri | 46.9491% |
| Picochlorum sp.SENEW3 | 60.2571% |
| Picochlorum costavermella | 59.5358% |
| Auxenochlorella protothecoides | 59.8031% |
| Chlorella variabilis NC64A | 61.1045% |
| Micractinium conductrix | 66.726% |
| Chlorella sp.Dachan | 72.5386% |
| Chlorella sorokiniana BD08 | 71.5315% |
| Chlorella sorokiniana BD09 | 73.3217% |
| Chlorella sorokiniana UTEX 1602 | 100% |

Figure S5. Identity comparation of amino-acid sequences of 14 algal genomes. a) A heatmap of average blast bit score of 14 genomes using the protein coding genes (e-value ≤ 1e-10). The self-blasted results were represented by the color grey. b) A bar chart of amino-acid similarity statistics of 13 genomes in comparation with *Chlorella sorokiniana* UTEX 1602.
